# A widespread role for SLC transmembrane transporters in resistance to cytotoxic drugs

**DOI:** 10.1101/726539

**Authors:** Enrico Girardi, Adrián César-Razquin, Konstantinos Papakostas, Sabrina Lindinger, Justyna Konecka, Jennifer Hemmerich, Stefanie Kickinger, Felix Kartnig, Alvaro Ingles-Prieto, Giuseppe Fiume, Anna Ringler, Charles-Hugues Lardeau, Richard Kumaran Kandasamy, Stefan Kubicek, Gerhard F. Ecker, Giulio Superti-Furga

## Abstract

The activity and potency of a drug is inherently affected by the metabolic state of its target cell. Solute Carriers (SLCs) represent the largest family of transmembrane transporters in humans and constitute major determinants of cellular metabolism. Several SLCs have been shown to be required for the uptake of individual chemical compounds into cellular systems, but systematic surveys of transporter-drug relationships in human cells are currently lacking. We performed a series of genetic screens in the haploid human cell line HAP1 using a set of 60 cytotoxic compounds representative of the chemical space populated by approved drugs. By using a SLC-focused CRISPR/Cas9 lentiviral library, we identified transporters whose absence induced resistance to the drugs tested. Among the hundreds of drug-SLC relationships identified, we confirmed the role of the folate transporter SLC19A1 on the activity of antifolates and of SLC29A1 on several nucleoside analogs. Among the newly discovered dependencies, we identified the transporters SLC11A2/SLC16A1 for artemisinin derivatives and SLC35A2/SLC38A5 for cisplatin. The functional dependence on SLCs observed for a significant proportion of the compounds screened suggested a widespread role for SLCs in the uptake and cellular activity of cytotoxic drugs and provided an experimentally validated set of SLC-drug associations for a number of clinically relevant compounds.

## Introduction

Cellular metabolism influences the rates of drug uptake and extrusion/excretion through the action of transmembrane transporters, the availability of cofactors and target(s) and the processing of prodrugs into active forms^1^. Moreover, drug modifying enzymes (DMEs), such as members of the cytochrome C family and glucosyltransferases, add functional groups to xenobiotic compounds, facilitating their removal from the cell and eventually the organism^1^. Most of what is known about the uptake of drugs by membrane-bound transporters stems from the analysis of drug disposition in the kidney, liver, intestine and blood-brain-barrier, with a particular focus on the entry and exit of pharmacological agents from the blood circulation^2,3^. In particular, two main families of transporters have been previously shown to directly interact with drugs: ATP-binding cassette transporters (ABCs)^4^ and Solute Carriers proteins (SLCs)^5^. ABC transporters are generally involved with the export of drugs, while SLCs have been mostly described to be involved in compound uptake even though exceptions to this rule exist, such as in the case of the MATE (multidrug and toxic compound extrusion) transporters^6^. Notably, SLCs represent a largely understudied family, counting more than 400 members of which at least 30% are still considered entirely orphan^3^. SLCs are divided into subfamilies based on sequence similarity and have been shown to transport a variety of molecules, ranging from nucleotides, sugars and lipids to amino acids and peptides^3,5^, often with overlapping specificities. Consistent with their critical role in drug absorption and excretion, considerable knowledge has accumulated on a few large subfamilies of SLCs prevalently expressed in kidney, liver and intestine, such as the SLC22 and SLCO families^7,8^. There is ample consensus in ascribing an important role for these transporters in governing pharmacokinetics of several drugs, which has been corroborated by several pharmacogenomic polymorphisms^9^. Accordingly, the US Food and Drug Administration and the European Medicine Agency now recommend testing of several ABC and SLC22/SLCO members for clinical drug interaction studies^10^. However, it remains a matter of debate, to which extent membrane-bound transporters are involved in the uptake and metabolism of drugs at the target cell level, such as in muscle, brain or tumor cells^11–14^. Some drugs have been reported to depend on protein carriers to enter cells, with prominent cases such as the family of antifolate drugs (e.g. methotrexate, pralatrexate, raltitrexed) interacting with the folate transporters SLC19A1/RFC1 (reduced folate carrier 1) and SLC46A1/PCFT (proton-coupled folate transporter)^15^, or the nucleoside transporter SLC29A1/ENT1 (equilibrative nucleoside transporter 1) interacting with several nucleoside analogs such as clofarabine, gemcitabine and fluorouracil^16^. In parallel, modulation of transporter activity or expression levels has been shown to affect the efficacy of drugs, independently from direct uptake events, through their effects on cellular metabolic processes such as glycolysis and oxidative phosphorylation^17–19^.

Genetic screening offers a powerful tool to identify both direct and indirect interactions between a gene and a specific phenotype. Using insertional mutagenesis, we recently demonstrated that the presence of the intact *SLC35F2* gene was the major determinant of the uptake of sepantronium bromide (YM155), a small molecule displaying anti-tumor activity *in vitro* and *in vivo*, in a variety of cell lines^20^. Similar forward genetics approaches have previously led to the identification of transporters involved in the uptake of cytotoxic compounds such as tunicamycin and 3-bromopyruvate^21,22^. Given the lack of molecular reagents available for the solute carrier family^3^, only recently have modern human cell genetic approaches allowed to test the hypothesis that SLC-mediated drug action, generally through uptake, is rather the rule than the exception. To tackle this important question in a focused way, we used an SLC-specific CRISPR/Cas9 KO library to perform a genetic survey of transporters involved with a chemically diverse set of 60 cytotoxic drugs. We identified and validated a large number of SLC-compound associations, providing insights into both direct uptake events and indirect associations affecting the metabolism and mechanism of action of the drugs tested.

## Results

### Generation of a SLC-specific CRISPR/Cas9 library

In order to investigate all SLC genes in an unbiased manner, we constructed a CRISPR/Cas9 library targeting 394 human SLC genes and pseudogenes with multiple single guide RNAs (sgRNAs) per gene. Particular care was taken to avoid sgRNAs with sequences sharing similarity with other SLC or ABC transporters. A set of negative control sgRNAs (predicted not to target any sequence in the genome) as well as a set of sgRNAs targeting genes scoring as essential in the HAP1 and KBM7 cell lines, based on previous insertional mutagenesis data^23^, were also included in the pool (Fig 1a, Suppl Table 1). The resulting library consisted of 2,609 unique sgRNAs, allowing for highly scalable and multiplexable screening and sequencing protocols. Presence of all sgRNAs was confirmed by Next Generation Sequencing (NGS, Suppl Fig 1a). Comparison of plasmid samples with HAP1 cell genomic DNA samples taken nine days post-infection showed significant depletion of sgRNAs targeting the set of essential genes (54/120, p-value = 8.2 × 10^-26^, Fisher’s exact test, Suppl Fig 1b). No significant depletion or enrichment was observed for the set of negative control sgRNAs (21/120, enrichment p-value = 0.29, depletion p-value = 1, Fisher’s exact test, Suppl Fig 1c). At the gene level, we identified several SLCs important for optimal fitness of HAP1 cells, including SLC35B1, the recently deorphanized ATP/ADP exchanger in the endoplasmic reticulum^24^ and MTCH2, a mitochondrial carrier involved in the regulation of apoptosis^25^ (Suppl Fig 1d). To validate the efficiency and specificity of our library in detecting SLCs associated with drug action, we screened for SLC genes responsible for resistance to YM155. Screening in HAP1 cells with 200nM YM155 for 72h resulted in a clear enrichment in sgRNAs targeting the SLC35F2 gene (Suppl Fig 1e), confirming that SLC35F2 is the sole SLC responsible for YM155 resistance and consistent with our previous results derived from insertional mutagenesis experiments^20^.

**Figure 1.**
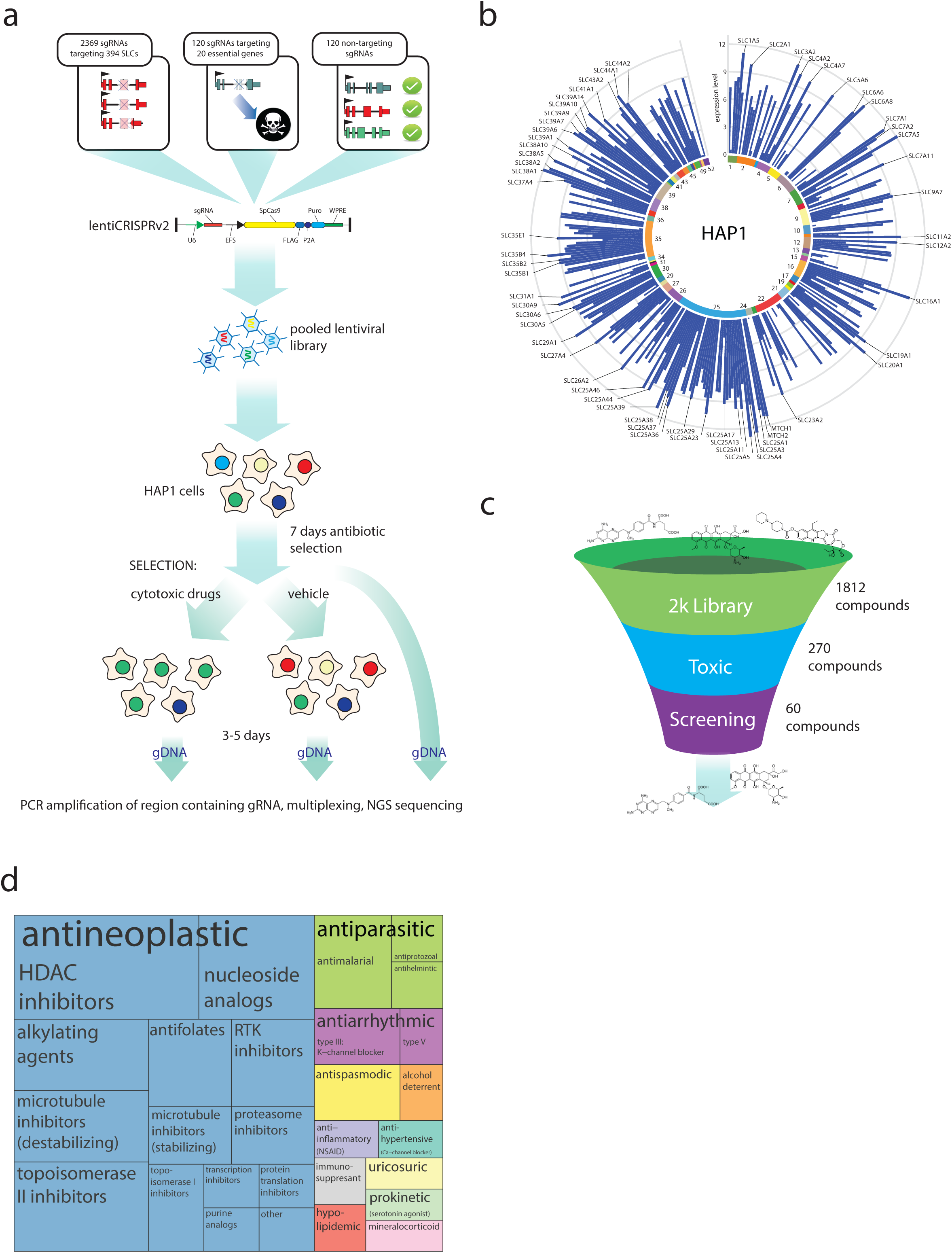
**a.** Schematic view of the composition of the SLC-focused CRISPR/Cas9 library and experimental outline of the genetic screen. **b.** Circular plot showing SLCs expressed in HAP1 cells according to RNAseq data from Brockmann *et al*. SLC families are indicated in the inner circle while transcript expression level (log_2_ counts per 10^7^ reads) is shown as blue bars. SLCs with an expression level above 9 are labeled. **c.** Schematic view of the compound sets and steps applied to the selection of a final set of drugs for screening. **d.** Treeplot view of the drug classes and subclasses included in the screening set.

### The SLC repertoire of HAP1 cells

Immortalized human cell lines typically express 150-250 SLC genes, with abundancy patterns resembling those of tissues^26,27^. For our screen, we chose HAP1 cells, a human cell line bearing considerable technical advantages^28,29^. HAP1 cells express 207 human SLC genes, as assessed by transcriptional profiling using RNA-Seq^30^ (Fig 1b). Importantly, these cells do not express most members of the organic ion transporter SLC22 family that have been implicated in the uptake of drugs in kidney, gut and liver^7^, making it ideally suited to test the potential role of other SLC families. Moreover, by being haploid, loss-of-function phenotypes induced by CRISPR/Cas9 technology should be more easily interpretable, as they do not represent composite mutants of different alleles.

### Identification of a set of cytotoxic drugs

For our genetic screens, we aimed at selecting a set of compounds representative of the chemical and functional space populated by drugs. We therefore tested cytotoxicity of a set of 1812 compounds (2k library) including the CLOUD library^31^ and the NIH Clinical Collection as well as sets of epigenetic modifiers and toxic compounds. A subset of 270 (14.9%) compounds was found to be cytotoxic in HAP1 cells at the tested concentration (toxic set) (Fig 1c). A eight-point dose-response curve was subsequently performed for each compound to determine IC_50_ values, after including an additional set of drugs with underrepresented indications e.g. compounds involved in DNA-damage-based-sensitivity. A group of 60 compounds chosen to cover different target classes by focusing on investigational/approved drugs with clinical relevance and diverse indications was finally selected for screening with the CRISPR/Cas9 library (screen set, Fig 1d, Suppl Table 2).

### Genetic screening identifies known and novel SLC-drug associations

We infected haploid HAP1 cells with the SLC CRISPR/Cas9 library to generate a pool of cells each lacking one specific SLC. The population was treated with multiple concentrations, generally one, three and ten times the measured IC_50_, of the cytotoxic compounds for 72h. As expected by dosing cytotoxic compounds, we retrieved all the samples treated with the IC_50_ concentrations as well as 78% (35/45) of the treatments at 2-3X the IC_50_ and 37% (22/60) of the 10X IC_50_ treatments. Enrichment was first calculated at the sgRNA level using DESeq2^32^ (Fig 2a) and then aggregated at the gene level using the GSEA algorithm^33^ (Fig 2b). When positive enrichment for SLC genes was calculated, we identified 201 SLC/drug associations involving 47 drugs (76 different treatment modalities) and 101 SLCs (Figure 2c, Suppl Fig 2a-b, Suppl Table 6) at a False Discovery Rate (FDR) ≤1%. Most of the SLCs identified are expressed in HAP1 cells (93/101, 92%, Suppl Fig 2c), with a large proportion localized at the plasma membrane (Suppl Fig 2d).

**Figure 2.**
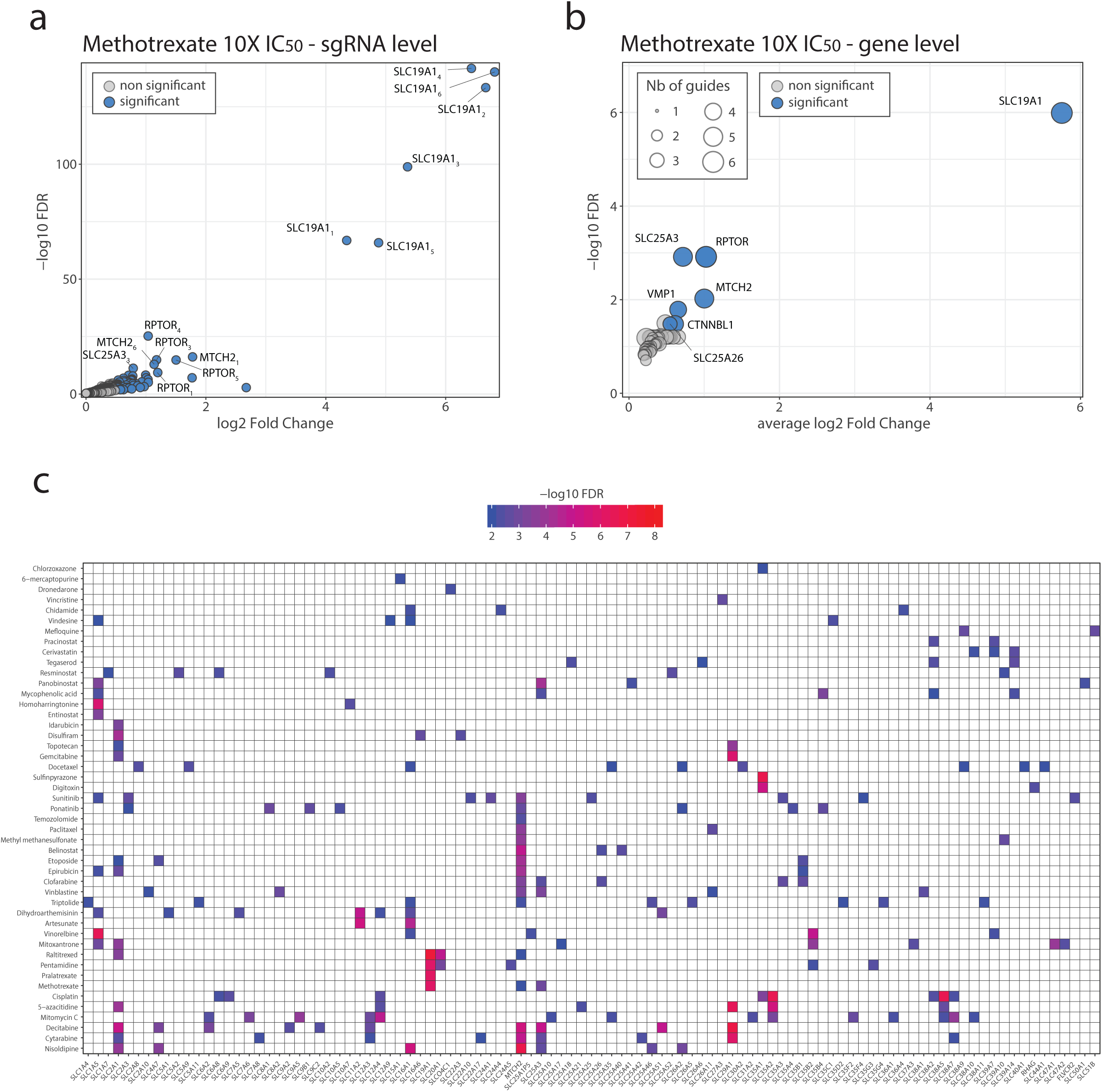
**a.** sgRNA-level enrichment for samples treated with 10X IC_50_ methotrexate, as determined by DESeq2. All six sgRNAs targeting the SLC19A1 gene showed significant enrichment. **b.** Gene-level enrichment for samples treated with 10X IC_50_ methotrexate, as determined by GSEA. Average log2 fold change for the significant sgRNAs for each gene is shown in the x-axis. Circle size indicates the number of significant sgRNAs. **c.** Overview of significantly enriched SLCs (FDR≤1%) identified upon treatment with different compounds. Significant enrichments for all different doses of the same compound are merged together (union), always selecting the most significant value for repeated hits. SLC genes are ordered by name, and treatments are ordered by hierarchical clustering based on the gene-level results. Results are derived by pooling data from at least two independent experiments.

Rewardingly, drugs belonging to the same classes generated prominent clusters. One example is represented by the cluster of the antifolate drugs methotrexate, raltitrexed and pralatrexate, which all induced a strong enrichment in KOs of the reduced folate carrier SLC19A1/RFC. This transporter has been previously recognized as the main uptake route of these antimetabolites into cells^15^. In particular, pralatrexate was developed to exploit this entry route^34^ and it showed exclusive enrichment for SLC19A1 in our screen (Fig 2c). Interestingly, within this cluster we found the structurally unrelated drug pentamidine, which is used for the treatment of African trypanosomiasis and leishmaniasis, as well as for the prevention and treatment of pneumocystis pneumonia (PCP) in immunocompromised patients. The mechanism of action (MoA) of this drug is poorly understood but earlier reports suggested it might be involved with inhibition of the parasite dihydrofolate reductase^35^. Another cluster included the nucleoside-like drugs decitabine, cytarabine, 5-azacytidine and gemcitabine, which all showed enrichment for the nucleoside transporter SLC29A1/ENT1 (Fig 2c). SLC29A1 was previously reported to act as an importer of these compounds^16,36,37^. As HAP1 cells express very low levels or do not express the additional nucleoside transporters SLC29A2, SLC28A1 and SLC28A3 (Fig1b), loss of SLC29A1 is expected to result in an impaired uptake of these compounds within the cell. Interestingly for some (i.e. cytarabine and decitabine), but not all of these compounds, we detected enrichment of the mitochondrial phosphate transporter SLC25A3 (Fig 2c).

We also observed more exclusive interactions, such as the one between the antineoplastic drug mitoxantrone and the two transporters MATE1/SLC47A1 and MATE2/SLC47A2 (Fig 2c). While mitoxantrone, a type II topoisomerase inhibitor and DNA intercalating agent, was previously reported to also inhibit the uptake/efflux of MATE1/SLC47A1 substrates^38,39^, our findings suggest that the interaction of this compound with these transporters may be associated with their uptake.

Moreover, we observed several cases of interactions offering plausible insights in the MoA or metabolic impact of a drug treatment. The artemisinin-derivatives artesunate and dihydroartemisinin showed an enrichment for the SLC11A2 and SLC16A1 genes (Fig 2c). These compounds, generally used for the treatment of malarial infections, have recently found additional use as antineoplastic agents^40,41^. Although the MoA is not fully understood, their cytotoxicity appears to rely on an iron/heme-dependent activation step and subsequent generation of Reactive Oxygen species (ROS)^42^. SLC11A2, also known as DMT1 (divalent metal transporter 1), is a metal transporter which has been shown to control the pool of cytoplasmic iron^43^. SLC16A1, also known as MCT1 (monocarboxylate transporter 1) is a major lactate exporter that plays an important role in glycolytic metabolism^44^ and could be directly involved in drug uptake or affect the ROS response to these compounds. Finally, we also observed a very strong enrichment of the transporters SLC35A2, a nucleoside-sugar Golgi transporter^45^, and SLC38A5^46^, an amino acid transporter, upon treatment with the DNA-damaging agent cisplatin (Fig 2c). Overall, the experimental drug-SLC gene interaction map showed a remarkably large landscape of known and novel associations covering 35 different SLC subfamilies, representing almost two thirds of the total of subfamilies tested.

### Validation of selected SLC-drug associations by Multicolor Competition Assay

While the screen was effective in determining a genetically defined functional relationship, it did not reveal the degree and the kinetics by which loss of function of an individual SLC affected loss of cell growth compared to an isogenic cell. We selected a set of 34 SLC-drug interactions (Suppl Table 4), involving 21 drugs and 13 SLCs, to assess growth differences in pairwise comparisons. We applied a FACS-based Multicolor Competition Assay (MCA), an approach that has been previously used to validate forward genetics screen results^47^. In this assay, HAP1 cells carrying a sgRNA targeting a given SLC (Suppl Table 5) and an enhanced GFP (eGFP) expression construct were mixed at 1:1 ratio with cells carrying a control sgRNA (targeting the *Renilla spp luciferase* gene, not present in the cells) and a mCherry construct. The mixed population was then treated with either vehicle or the cytotoxic compound at 1-3 times the IC_50_ and the ratio of GFP+/mCherry+ determined by FACS three and ten days later (Fig 3a). Two sgRNAs were used for each gene targeted in order to control for sgRNA-specific effects. Overall, this approach enabled the validation of several among the strongest interactions derived from the genetic screen (Fig 3b, Suppl Table 7). In particular, we confirmed the strong effects of SLC19A1 and SLC29A1 loss on the resistance to antifolate and nucleoside analogs at both early (3 days) and late (10 days) timepoints. In addition, we validated the effect of the loss of SLC20A1, a phosphate transporter, upon methotrexate treatment. We also observed strong and time-dependent enrichments of cells lacking SLC11A2 or SLC16A1 upon treatment with artesunate and dihydroartemisinin, as well as in the case of panobinostat and the amino acid transporter SLC1A5. Finally, we observed strong enrichments of SLC35A2- and SLC38A5-lacking cells upon cisplatin treatment. This effect was already discernible for SLC38A5 after three days of drug exposure and became clear for both genes after ten days. Overall, we were able to confirm the majority (26/34) of the associations tested at one or more timepoints, therefore validating the approach and results of the genetic screen.

**Figure 3:**
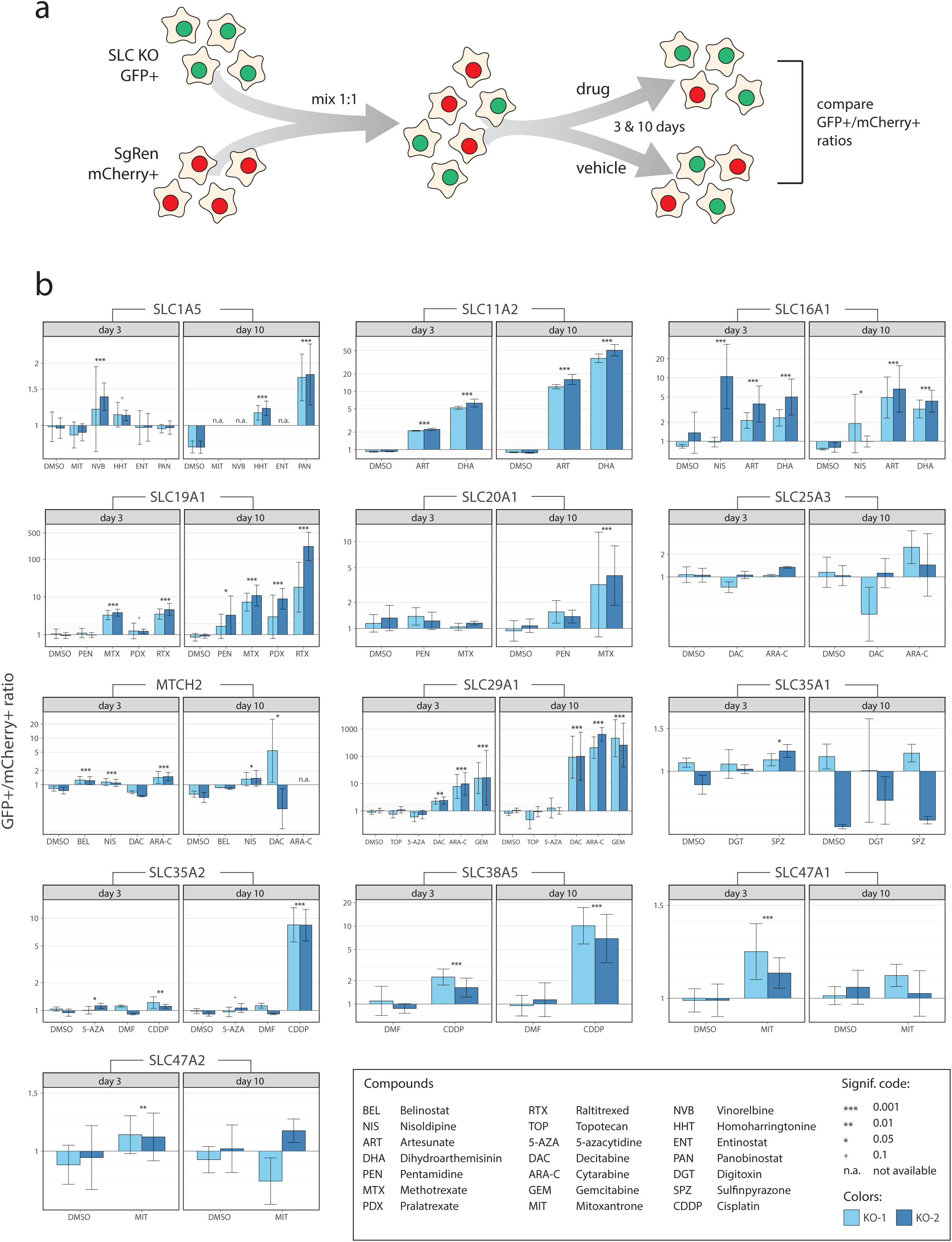
**a.** Schematic view of the Multicolor Competition Assay (MCA). **b.** Validation of selected SLC/drug associations by MCA. Results are shown by gene tested, pooling data of at least two independent experiments each performed in technical triplicates. Ratios of GFP+/mCherry+ populations normalized to the day0 ratios are shown for the indicated SLC/drug combinations at the given timepoints as mean(log(ratios)) ± sd(log(ratios)). Statistical significance was calculated by ANOVA followed by Dunnett’s test. n.a. denotes cases where no live cells were measured.

### SLC16A1 protein levels affect sensitivity to artesunate

We further validated the interaction between artesunate (Fig 4a) and SLC16A1 as an example of novel SLC-drug association. To further confirm this interaction, we made use of two single cell-derived HAP1 cell lines carrying frameshift mutations in SLC16A1 (ΔSLC16A1_1, ΔSLC16A1_2). These cell lines showed increased resistance to artesunate treatment compared to WT cells, as assessed by a luminescence-based cell viability assay (Fig 4b). Importantly, ectopic expression of a SLC16A1 cDNA resulted in increased sensitivity to artesunate, compared to control cell lines ectopically expressing eGFP (Fig 4c-d) demonstrating that SLC16A1 protein levels affect cell sensitivity to artesunate.

**Figure 4:**
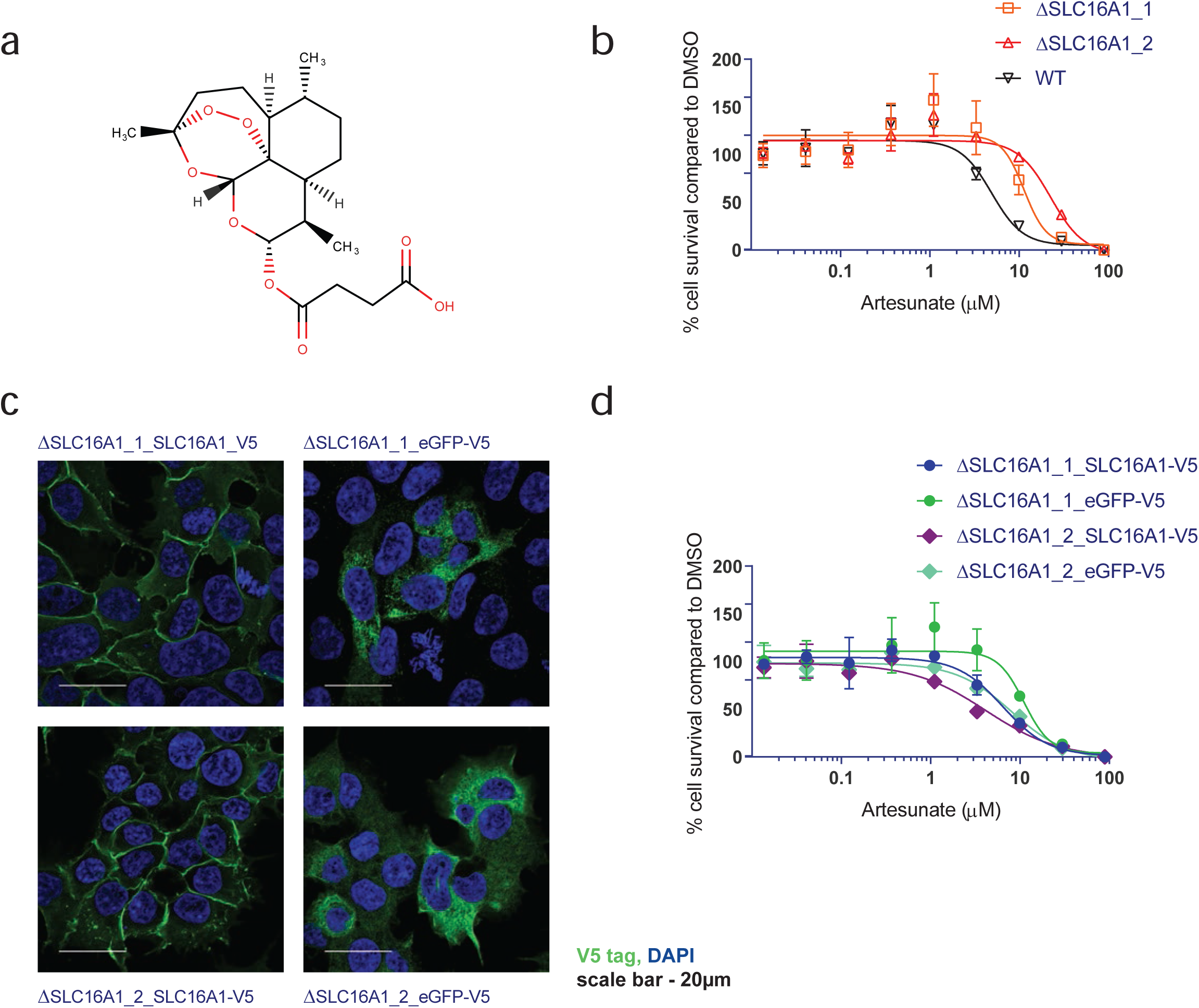
**a.** Chemical structure of artesunate **b.** Cell viability assay comparing sensitivity to artesunate of WT HAP1 cells and cells carrying frameshift mutations in the SLC16A1 gene (ΔSLC16A1_1, ΔSLC16A1_1). **c.** Cell viability assay showing increased sensitivity of HAP1 SLC16A1 KO cells reconstituted with SLC16A1 cDNA compared to cells reconstituted with eGFP. **d.** Confocal images of HAP1 cells lacking endogenous SLC16A1 and reconstituted with GFP or SLC16A1 cDNA. Green: eGFP or SLC16A1, Blue: DAPI. Scale bar: 20 μm.

### The screened and SLC-associated compounds sets are representative of the known chemical space

The remarkable finding that so many drugs showed a functional dependence on an SLC transporter raised the question whether the functional landscape tested is biased for particularly large and hydrophilic compound and not representative of the general drug-like chemical space. In order to assess if the selected set of compounds (screen set) was a representative subset, we performed a detailed cheminformatics analysis. Drugbank 5.1.1^48^ was used as a reference of the known drug chemical space and compared to all three aforementioned compound sets (2k library, toxic and screen, Fig 1c, Fig 5a, Suppl Fig 3a). All sets were curated according to the same protocol (see methods) and 22 physicochemical 2D descriptors (Suppl Table 3) were calculated for every compound. Comparison of individual descriptor mean and median values showed no strong bias across the four compound sets (Suppl Fig 4). In order to visualize the distribution of all compounds in the chemical space, a principal component (PC) analysis was performed. The first and the second PCs were able to explain 62.1 % of the variance of the data (Fig 5a-b). Descriptors contributing the most to the variance of PC1-2 were number of heavy atoms, molecular weight, Labute’s surface area, number of heteroatoms, number of saturated rings, number of H-bond donors and polar surface area (TPSA, Fig 5c). Importantly, compounds of all sets were similarly distributed along the two first PCs. Overall, this analysis showed that there is no striking difference in the distribution of physicochemical properties of the compound sets used in this study. Therefore, the final screen set can be considered representative of the general drug chemical space. Moreover, we compared the chemical properties of the set of 47 compounds associated to at least one SLC (active) and the compounds with no associations (inactive) to the DrugBank dataset (Fig 5d, Suppl Fig 3b) as well as between them (Suppl Fig 3c) and observed no trend that would suggest a presence of specific properties in the set of drugs showing associations with SLCs.

**Figure 5:**
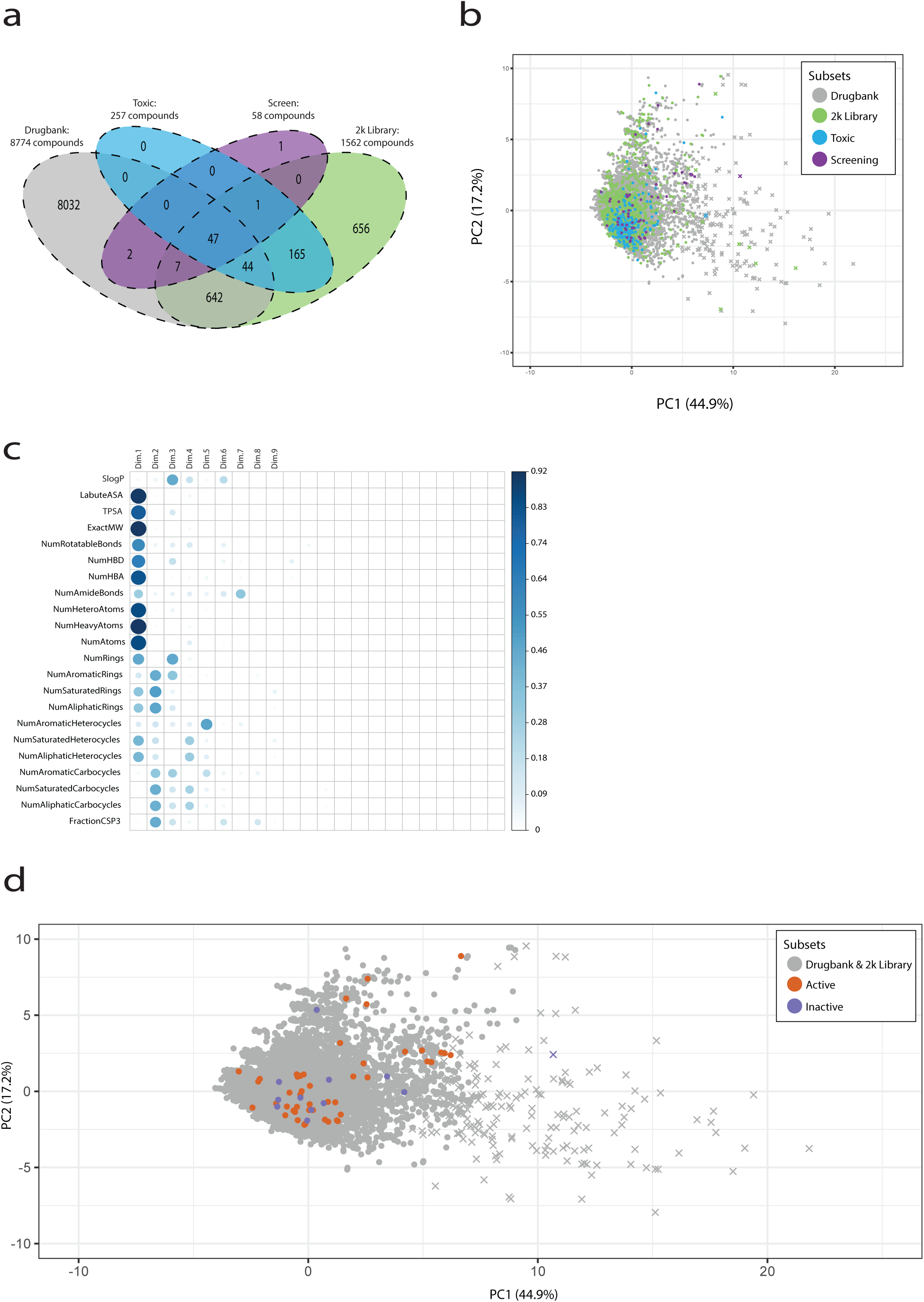
**a.** Venn diagram showing the compound subsets used for the chemoinformatic analysis after stripping of stereochemistry and removal of anorganic and duplicated compounds. **b.** Principal component analysis of compounds in the DrugBank set of reference as well as in the sets tested in this study based on 22 annotated 2D chemical descriptors. Zoomed-in ssssssssversion for clarity, the full plot is shown in Suppl Fig 3a. Compounds with a molecular weight below 900 Da (defined as “small molecule” by DrugBank) are shown as circles, the remaining compounds as crosses. **c.** Correlogram plot showing the 2D descriptors contribution to the PCA analysis. **d.** Principal component analysis of compounds in the DrugBank set of reference compared the SLC-associated (active) and non-SLC-associated (inactive) compounds based on 22 annotated 2D chemical descriptors. Zoomed-in version for clarity, the full plot is shown in Suppl Fig 3b.

## Discussion

Transmembrane transporters represent a major class of metabolic genes involved in several cellular processes affecting drug potency and activity, including the uptake and extrusion of these xenobiotic compounds^2^. In the past, the discovery of specific transporters for several cytotoxic compounds by insertional mutagenesis and CRISPR/Cas9-based screens had provided clear examples of the power of genetic approaches for the identification of such relationships^20–22,49^. However, despite the unambiguous involvement of these transporters in drug uptake, it is possible that these relationships are exceptional in nature and confined to particular chemical subtypes. As an alternative possibility, most drugs thought to act on an intracellular target would indeed require a membrane-spanning transporter to gain access to the inside of cells. The reasons why these had not yet been identified could have been the lack of convenient genetic tools in cellular intact systems. As motivation for this study, we reasoned that a focused forward genetic approach in live human cells (Fig 1a, Suppl Table 1) would allow us to systematically investigate the frequency by which a drug or drug-like compound would be affected by the function of an SLC gene.

As read-out compatible with genetic screening, we opted for simple cellular survival, as it allows to monitor strong selective pressures and to focus on cytotoxic/cytostatic compounds of clinical relevance. A cheminformatics analysis of the set of 60 screened compounds, including several approved drugs (Fig 5a-c, Suppl Table 2), showed no bias in physicochemical properties compared to the DrugBank database, thus supporting its use as a set representative of the chemical space occupied by drugs.

Antimetabolites such as antifolates and nucleoside-analogs scored strongly in our setting, recapitulating the known cases of drug uptake mediated by transporters such as SLC19A1 and SLC29A1 (Fig 2c, 3b). Interestingly, we also identified several additional strong interactions across different drug classes, such as the role of the iron transporter SLC11A2 in determining resistance to artemisinin derivatives (Fig 2c, 3b). This is consistent with the correlation between intracellular iron levels and drug cytotoxicity previously suggested for these compounds^50^. We also validated interactions between artesunate/dihydroartemisinin and the monocarboxylate transporter SLC16A1, showing in the latter case that SLC16A1 protein levels determine the sensitivity to artesunate (Fig 4), as well as between cisplatin and the transporters SLC35A2 and SLC38A5. The latter is particularly interesting as SLC38A5 is a glutamine transporter expressed at high levels in cells of hematopoietic origin and several studies reported a dependence on glutaminolysis for cisplatin-resistant cells^51,52^. Interestingly, we observed several interactions comprising key, often essential, transporters involved in major energetic pathways such as SLC2A1/GLUT1, the major glucose transporter at the plasma membrane, SLC25A3, the mitochondrial phosphate transporter, or MTCH2, a mitochondrial carrier involved in apoptosis regulation (Fig 2a, 3b). It has been shown that resistance to cytotoxic drugs often requires major metabolic rearrangements: e.g. glutaminolysis and cisplatin resistance^51,52^, a switch to oxidative phosphorylation in cytarabine resistance^53^ or drug-specific dependence on glycolysis^17^. The fact that several of the SLC-drug associations identified involve SLCs important for cellular fitness therefore speaks to the enormous metabolic pressure a cytotoxic drug imposes on a target cell.

Importantly, almost 80% (47/60) of the small chemical molecules tested were functionally dependent on an SLC gene (Fig 2c). The large number and proportion of novel drug-SLC relationships identified here strongly argues for a more general role of transporters than currently appreciated in the uptake and activity of drugs in target tissues. Previous surveys of cytotoxic compounds in yeast showed that transporter deletion affected the activity and uptake of ∼70% of the compounds tested (18/26), a proportion remarkably similar to the one observed in this study^54^. In light of the outcome of these systematic functional surveys, the notion that membrane permeability and bioavailability of drugs primarily reside in their ability to diffuse across membranes may require revision. As for the remaining 20% of compounds that did not show an association with SLCs, we did not observe any striking difference in their chemical properties when compared to the SLC-associated ones (Fig 5d), suggesting that these molecules may exert their activity or access cells through SLCs with redundant function. In these scenarios, genetic depletion of a single SLC would not be sufficient to score in our experimental set-up and higher-order genetic perturbations, of the type that could be achieved with vectors bearing multiple sgRNAs^55,56^, may be required. It is also possible that proteins other than SLCs are involved in drug uptake, such as yet poorly characterized TMEM proteins^57^, ion channels or entirely uncharacterized proteins. Larger focused libraries or genome-wide CRISPR/Cas9 screens may therefore reveal the involvement of non-redundant and non-essential proteins exerting drug-transporting function. In any case, it is now feasible and urgent to investigate the genetic determinants of drug activity and especially uptake. This first systematic functional survey in intact human cells seems to severely disavow the hypothesis, still central to most medicinal chemistry and many biochemistry textbooks, that given the proper physicochemical parameters, chemical entities will be able to enter cells by diffusion^58^. The evidence provided here is clearly beyond anecdotal and will hopefully trigger further campaigns of similar scope. Knowledge of the transporters affecting uptake and activity of drugs in tumors and tissues is certain to represent a cornerstone of precision therapy of the future. Moreover, the relationship between the expression of SLCs, cellular/organismal metabolism and nutrition is likely to allow the opening of additional therapeutic windows.

## Materials and methods

### Generation of a SLC-wide CRISPR/Cas9 lentiviral library

A set of single guide RNAs (sgRNAs) targeting 388 human SLC genes, generally with six sgRNAs per gene, were manually selected (or generated) to include sequences with predicted high efficiency and specificity, as assessed in Doench *et al*^59^, and to minimize targeting of other SLCs or of ABC transporters (Suppl Table 1). sgRNAs targeting six SLC pseudogenes (SLC7A5P1, SLC7A5P2, SLC9A7P1, SLC2A3P1, SLC25AP5, SLC35E1P1) for which transcription was previously reported in at least two expression datasets (FANTOM5, CCLE, ENCODE, Cosmic, GENCODE, Uhlen et al, Illumina)^60–66^ were also included. An additional set of 120 sgRNAs targeting 20 genes essential in both KBM7 and HAP1 cells^23^ based on the number of retroviral insertions observed were also selected (Suppl Table 1). Finally, a set of 120 non-targeting sgRNAs was included by generating random 20-mers and selecting for sequences with at least three (for the strong PAM NGG) or two (for the PAM NAG) mismatches from any genomic sequence with E-CRISP Evaluation^67^. Adapter sequences were added to the 5’ and 3’ sequences (5’prefix: TGGAAAGGACGAAACACCG, 3’suffix: GTTTTAGAGCTAGAAATAGCAAGTTAAAATAAGGC) to allow cloning by Gibson assembly in the lentiCRISPRv2 vector (Addgene #52961). The oligos were synthetized as a pool by LC Sciences. Full-length oligonucleotides (74 nt) were amplified by PCR using Phusion HS Flex (NEB) and size-selected using a 2% agarose gel (Primers: SLC_ArrayF TAACTTGAAAGTATTTCGATTTCTTGGCTTTATATATCTTGTGGAAAGGAC GAAACACCG, SLC_ArrayR ACTTTTTCAAGTTGATAACGGACTAGCCTTATTTTAACTTGCTATTTCT AGCTCTAAAAC)

The vector was digested with BsmBI (NEB) for 1h at 55°C, heat inactivated for 20’ at 80°C, following by incubation with Antarctic phosphatase (NEB) for 30’ at 37°C. A 10 μl Gibson ligation reaction (NEB) was performed using 5 ng of the gel-purified inserts and 12.5 ng of the vector, incubated for 1h at 50°C and dialyzed against water for 30’ at RT. The reaction was then transformed in Lucigen Endura cells and plated on two 245 mm plates. Colonies (equivalent to approximately 200X coverage) were grown at 32°C for 16-20h hours and then scraped from the plates. The plasmid was purified with the Endo-Free Mega prep kit (Qiagen). Of note, the library generated as described here, has been already successfully used in focused screens for phagocytosis^68^, necroptosis^69^ and cell survival upon viral infection^70^.

### Library NGS sequencing

Initial amplification of the library for NGS sequencing was performed by a two-step PCR protocol as described in Sanjana *et al* ^71^. Due to the presence of unspecific bands affecting the quality of the sequencing experiments, later samples were processed with a single-step PCR derived from Konermann *et al* ^72^. The PCR primers used to add barcodes and Illumina adapters were modified to allow for double indexing of samples.

### Enrichment analysis

sgRNA sequences were extracted from NGS reads, matched against the original sgRNA library index and counted using an in-house python script. Samples with less than 10^5^ total reads were excluded from further analysis. A two-step approach was implemented in order to obtain a final list of enriched candidate genes. First, differential abundance of individual sgRNAs was estimated using DESeq2 v1.20^32^. Models accounted for both treatment and time variables when time 0 samples were available; otherwise only the treatment factor was considered. Contrasts were performed individually for each treatment and dose vs controls (DMSO and untreated), and significance was tested using either one- or two-tailed Wald tests (i.e. alternative hypothesis LFC>0 for enrichment, and abs(LFC)>0 for enrichment or depletion, respectively). Then, sgRNAs were sorted by log2 fold change and aggregated into genes using Gene Set Enrichment Analysis (fgsea R package v1.7)^33,73^. To avoid false positives, only significant sgRNAs (p-value ≤ 0.05) were considered for enrichment, requiring also a minimum of two sgRNAs per gene. Gene enrichment significance was estimated by a permutation test using 10^8^ permutations, and p-values were corrected for multiple testing using the Benjamini-Hochberg procedure (FDR).

### Cell lines

HAP1 cells (Horizon Genomics) were grown in IMDM media (Gibco) supplemented with 10% FCS (Gibco) and 1% penicillin/streptomycin. For screening purposes, haploid cells were selected by FACS sorting after staining with Vybrant DyeCycle Ruby stain (Thermo Fisher Scientific), expanded for 3-5 days and frozen until further use. For CRISPR-based knockout cell lines, sgRNAs were designed using CHOPCHOP^71^ and cloned into pLentiCRISPRv2 (Addgene, #52961), LGPIG (pLentiGuide-PuroR-IRES-GFP) or LGPIC (pLentiGuide-PuroR-IRES-mCherry)^47^. sgRen targeting Renilla luciferase cDNA was used as negative control sgRNA^47^. Editing efficiency was determined with Tide-seq^74^. The SLC16A1 deficient clones (ΔSLC16A1_2882-2, renamed as ΔSLC16A1_2 in the text, and clone ΔSLC16A1_2882-10, renamed as ΔSLC16A1_1) were purchased from Horizon Genomics. Codon-optimized SLC16A1 cDNA or eGFP cDNA sequences were obtained from the ReSOLUTE consortium (www.re-solute.eu) and cloned in the pLX304 vector (Addgene plasmid #25890).

### Drug cytotoxicity screens

To mimic the genetic screen conditions, HAP1 cells were infected with a lentiCRISPRv2 vector carrying a sgRNA targeting the *Renilla* luciferase gene and selected with puromycin selection (1μg/ml) for 7 days. WT and lenti-infected cells were screened against a library composed of 1812 compounds at a single concentration in the range of 10-50μM. Viability was measured by CellTiterGlo assay (Promega) after 72h of treatment. DMSO and Digitoxin were used as negative and positive controls, respectively, to calculate cytotoxicity. Hits were defined as compounds giving more than 50% inhibition compared to DMSO controls. 8-point dose-response curves were performed to determine the IC_50_ values of the cytotoxic compounds in lentivirus-infected HAP1 cells.

### Chemical space analysis

Data curation was performed using a KNIME 3.6.0^75^ workflow which incorporates the python packages RDKit 2018.09.01^76^ and MolVS 0.1.1^77^ for handling and standardizing molecules (python 3.6.6^78^ was used). First, all compounds were neutralized by adding or removing protons. Then, compounds were cleaned by standardizing the representation of all aromatic rings, double bonds, hydrogens, tautomers and mesomers. Thereafter, all salts and mixtures were removed. In order to remove duplicates InChIKeys were calculated and all compounds were aggregated according to these InChIKeys. Chiral centers were also removed, as this stereochemistry information is often incorrectly assigned, which can lead to a lower detection rate of duplicates. Furthermore, only 2D descriptors were calculated, which cannot differentiate between enantiomers or diastereomers. It is noteworthy that the calculated 2D descriptors cannot be used to describe inorganic compounds, therefore those compounds were removed from further analysis. All 22 descriptors were computed with the RDKit nodes available in KNIME 3.6.0. Data visualization was performed in Rstudio 1.1.463^79^ with R 3.4.4^80^. Bar plots and violin plots were computed with ggplot2 3.1.0^81^, the correlation of descriptors plot (Suppl Fig 1g) was computed with corrplot 0.84^82^. Principal component analysis (PCA) was performed with the R packages factoextra^83^ and FactoMineR^84^.

### Genetic screens

Viral particles were generated by transient transfection of low passage, subconfluent HEK293T cells with the SLC-targeting library and packaging plasmids psPAX2, pMD2.G using PolyFect (Qiagen). After 24h the media was changed to fresh IMDM media supplemented with 10% FCS and antibiotics. The viral supernatant was collected after 48h, filtered and stored at -80°C until further use. The supernatant dilution necessary to infect haploid HAP1 cells at a MOI (multiplicity of infection) of 0.2-0.3 was determined by puromycin survival after transduction as described in Sanjana *et al*^85^. HAP1 cells were infected in duplicates with the SLC KO library at high coverage (1000x) and after selection for 7 days with puromycin (1 μg/ml) an initial sample was collected to control for library composition. Cells were then treated with multiple concentrations (generally 1X, 3X or 10X the IC_50_) of the cytotoxic compounds or vehicle (DMSO or DMF) controls for 72h and, when surviving cells were present, cell samples collected from both treated and control samples.

### Multicolor competition assay

Flow cytometry-based multi-color competition assays (MCA) were performed as described previously^47^. Briefly, HAP1 cells expressing LGPIC-sgRen (mCherry-positive) were mixed in 1:1 ratio with LGPIG (eGFP-positive) reporter cells containing sgRNAs targeting the gene of interest. The mixed cell populations were incubated with vehicle or drug for up to 10 days. The respective percentage of viable (FSC/SSC) mCherry-positive and eGFP-positive cells at the indicated time points was quantified by flow cytometry. Samples were analysed on an LSR Fortessa (BD Biosciences) and data analysis was performed using FlowJo software (Tree Star Inc., USA). Individual ratios were normalized to day 0 controls and then log transformed. In order to detect significant changes upon treatment, a two-way ANOVA model was fitted for every gene and day using treatment and KO as factors, and a one-tailed Dunnett’s test was performed to compare each treatment vs the control (DMSO for all drugs but cisplatin, DMF for cisplatin).

### Viability assays

For viability assays, 10000 HAP1 cells/well were plated in a 96-well plate and a 10-step, 3-fold dilution series performed in triplicates. Viability was measured by CellTiterGlo assay (Promega) after 72h of treatment.

### Confocal imaging

For the confocal imaging of 293T cells, high precision microscope cover glasses (Marienfeld) were coated with poly-L-lysine hydrobromide (p6282, Sigma-Aldrich) according to the manufacturers protocol. Cells were seeded onto cover glasses in normal growth medium and fixed in 4% Formaldehyde solution (AppliChem) in PBS 1x after 24 h of incubation. Permeabilization and blocking of samples was performed in blocking solution (10% FCS, 0.3% Saponin (47036, Sigma-Aldrich) in PBS 1x) for 1h rocking. Anti-V5 Tag primary antibody (Invitrogen, #46-0705) was diluted 1:500 in blocking solution and applied for 2h at room temperature, rocking. Samples were washed three times in blocking solution and anti-mouse Alexa Fluor 594 (Thermo, #A-11005) was applied 1:400 in blocking solution for 1h at room temperature, rocking. After three times washing in blocking solution nuclei were counterstained with DAPI 1:1000 in PBS 1x, for 10 min rocking. Cover glasses were mounted onto microscopy slides using ProLong Gold (Thermo Fischer Scientific) antifade mountant. Image acquisition was performed on a confocal laser scanning microscope (Zeiss LSM 780, Carl Zeiss AG), equipped with an Airyscan detector using ZEN black 2.3 (Carl Zeiss AG).

## Author contributions

E.G., G.S-F. conceived and designed the study. E.G., K.P., S.L., J.K., G.F., A.I. P., F.K., A.R. and C-H.L. performed experiments and analyzed data. A.C-R analyzed screening and validation data. J.H, S.Ki. and G.F.E. performed the chemoinformatic analysis. R.K.K. contributed to library design. S.Ku., G.F.E. and G.S-F. provided supervision.

## Acknowledgments

We thank all members of the Superti-Furga laboratory for discussions and feedback. We are also grateful to the Biomedical Sequencing facility for advice on Illumina sequencing and to the Flow Cytometry Core Facility of the Vienna Medical University for help with FACS sorting. We further thank Bojan Vilagos for graphical input and advice and Sara Sdelci for experimental advice. We acknowledge support by the Austrian Academy of Sciences, ERC grant to G.S-F. (ERC AdG 695214 GameofGates, E.G.), Austrian Science Fund (FWF I2192-B22 ERASE, A.C-R, FWF P29250-B30 VITRA, E.G, J.K, G.F,) and a Marie Sklodowska-Curie fellowship to E.G. (MSCA-IF-2014-661491). Research in the Kubicek laboratory is supported by the Austrian Federal Ministry for Digital and Economic Affairs and the National Foundation for Research, Technology, and Development, the Austrian Science Fund (FWF) F4701 and the European Research Council (ERC) under the European Union’s Horizon 2020 research and innovation programme (ERC-CoG-772437). The Pharmacoinformatics Reasearch Group (Ecker lab) acknowledges funding provided by the Austrian Science Fund FWF AW012321 MolTag.

