## Supplemental Figures 1-5 and Supplemental Tables 1-5 for "A widespread role for SLC transmembrane transporters in resistance to cytotoxic drugs"

**Supplemental Figure 1: a.** Violin plots of sgRNA count distributions in the SLC library plasmid samples and in the 9 days post-infection (p.i.) samples. **b.** Volcano plot (p-value vs. log2 fold change) for the differential representation of sgRNAs in samples collected 9 days p.i. vs. the plasmid library. P-values correspond to a two-tailed Wald test (DESeq2). sgRNA corresponding to the set of 20 essential control genes are shown in green. **c.** Same as b., but in this case sgRNAs corresponding to the set of 120 non-target negative control sequences are shown in red. **d.** Gene-level enrichment in 9 days p.i. vs. plasmid library. **e.** YM155 benchmarking screen. Read counts for the samples at day0, DMSO-treated and YM155-treated samples are shown.

**Suppl Figure 2: a.** Overview of significantly enriched SLCs (FDR≤1%) identified for drug treatments at different concentrations. SLC genes are ordered by name, and treatments are ordered by hierarchical clustering based on the gene-level results. **b.** Overview of significantly enriched SLCs (FDR≤1%) identified upon treatment with different compounds. Significant enrichments for all different doses of the same compound are merged together in order to ease interpretation (union), always selecting the most significant value for repeats. x and y-axis dendrograms display the hierarchical clustering of SLCs and treatments, respectively, calculated using the complete-linkage method with Euclidean distances based on gene-level adjusted p-values. **c.** Expression levels ($\log_2$ counts per $10^7$ reads) in HAP1 cells for SLCs significantly enriched in our screen. **d.** Localization of the SLCs significantly enriched in our screen. Data was assembled from the UniProt and Compartments databases followed by manual curation and annotation.

**Suppl Figure 3: a.** Example of the gating scheme used for the MCA assay. For this experiment, Hap1-Cas9 cells infected with lentiviral particles carrying sgRNAs targeting Renilla luciferase and either eGFP or mCherry fluorescent markers were mixed at 1:1 ratio and the relative abundance of the two populations assessed by FACS.

**Suppl Figure 4: a.** Principal component analysis of compounds in the DrugBank set of reference as well as in the sets tested in this study based on 22 annotated 2D chemical descriptors. Compounds with a molecular weight below 900 Da (defined as “small molecule” by DrugBank) are shown as circles, the remaining compounds as crosses. **b.** Principal component analysis of compounds in the DrugBank set of reference compared the SLC-associated (active) and non-SLC-associated (inactive) compounds based on 22 annotated 2D chemical descriptors. **c.** Principal component analysis of compounds in the SLC-associated (active) and non-SLC-associated (inactive) sets.

**Suppl Figure 5:** Plots of individual descriptor values across the four compound sets. Medians are indicated by triangles.

### Supplemental Figure 1

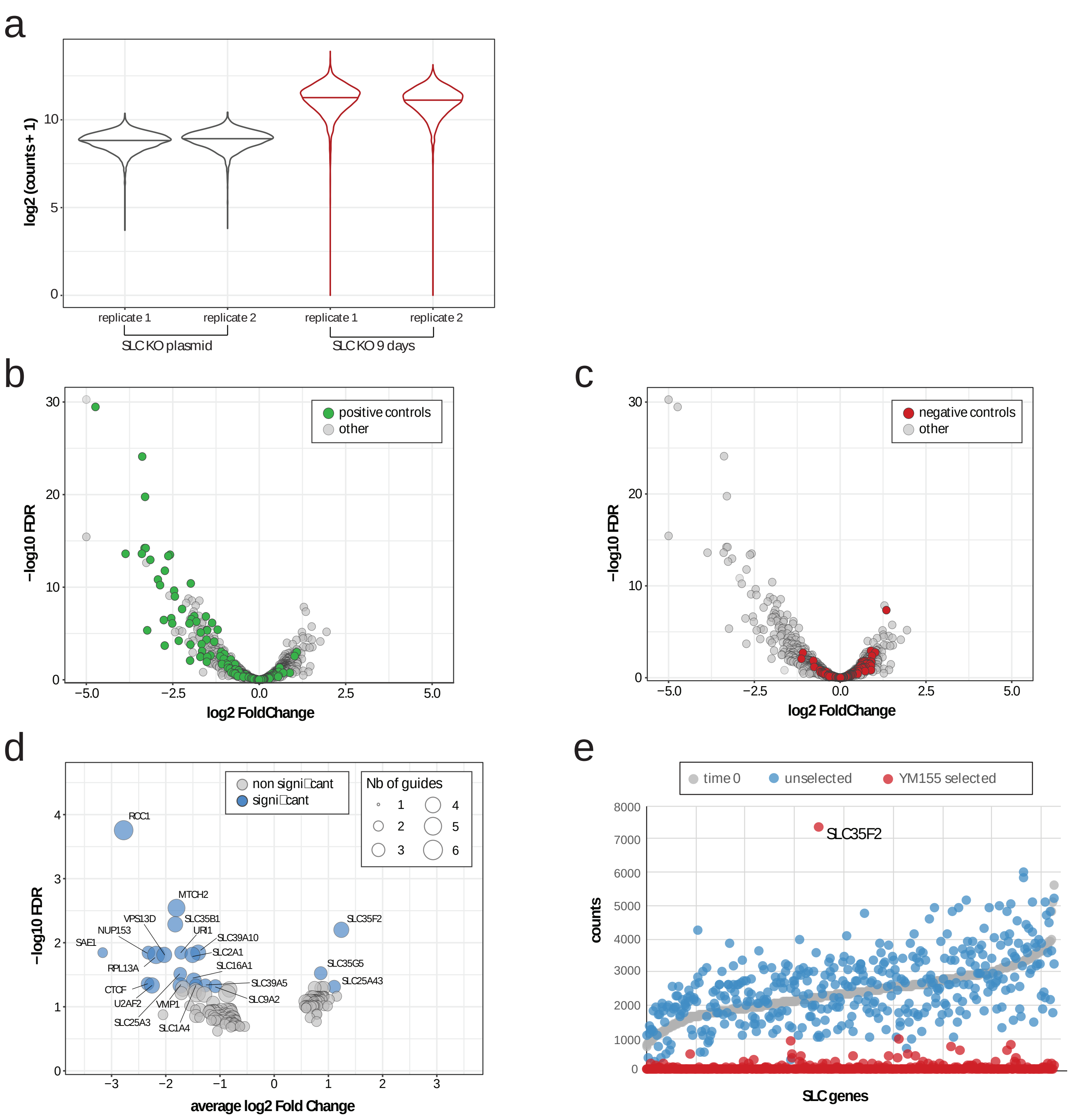

### Supplemental Figure 2

a

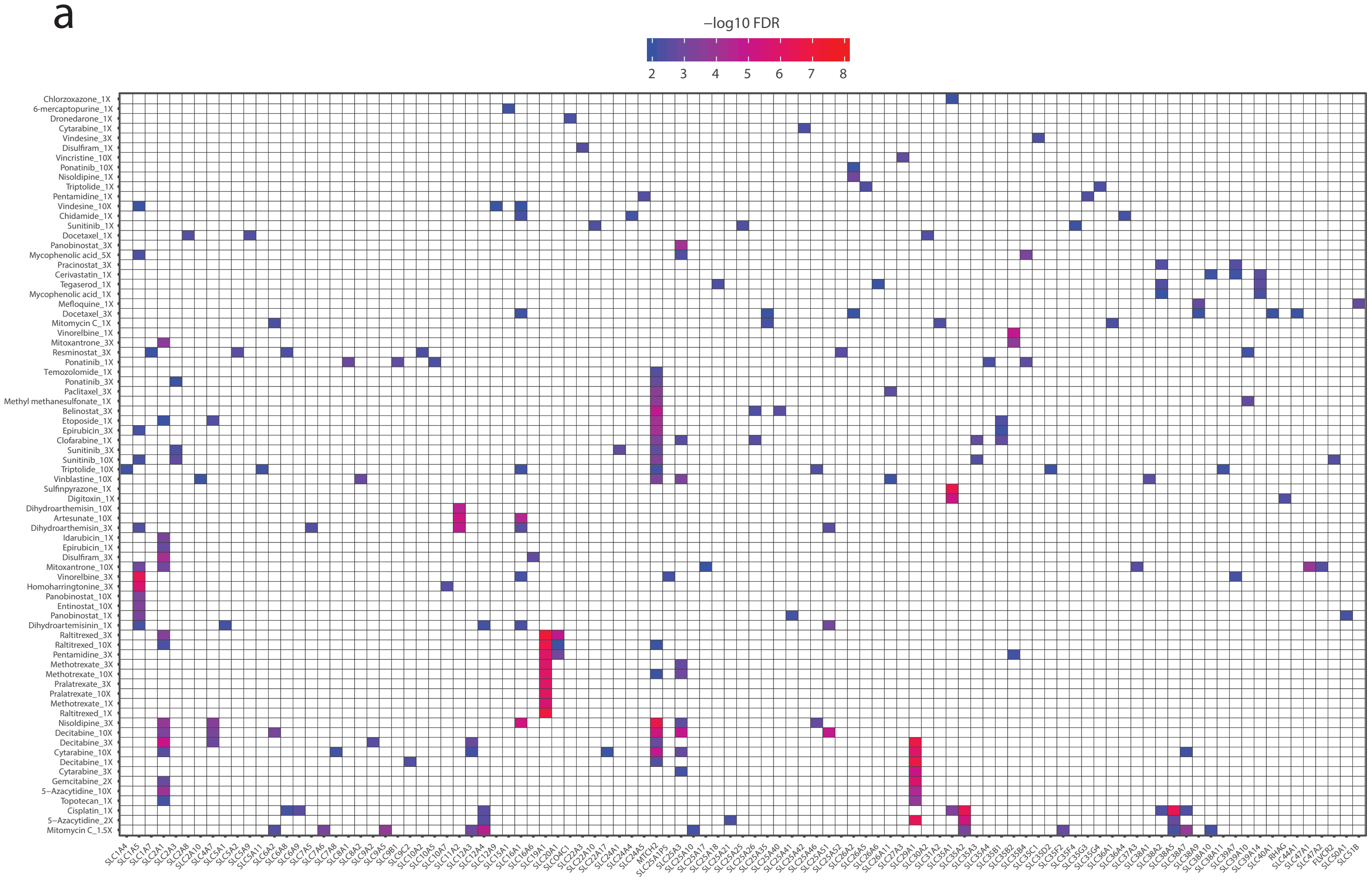


b

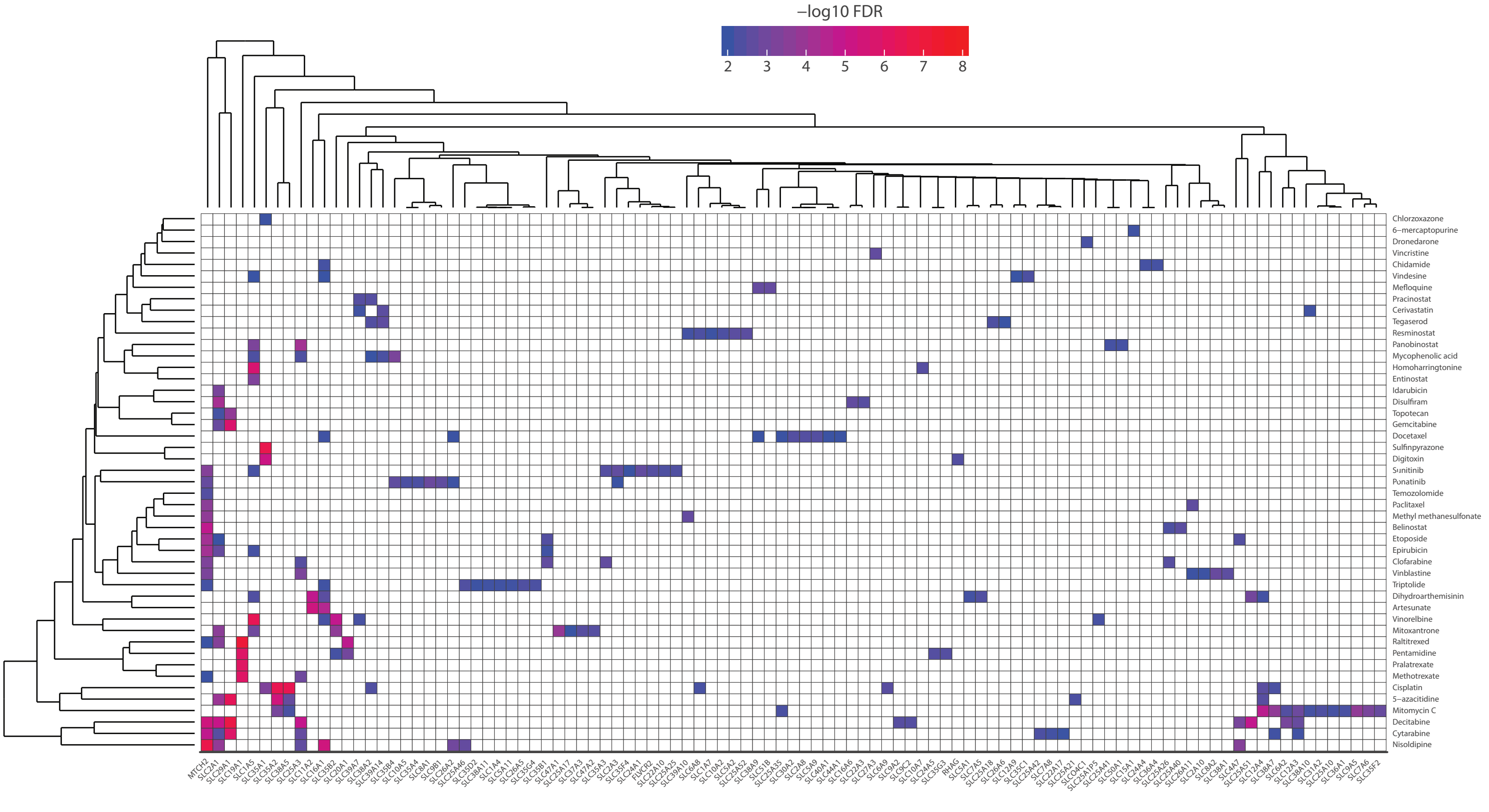

### Supplemental Figure 2 - continued

c

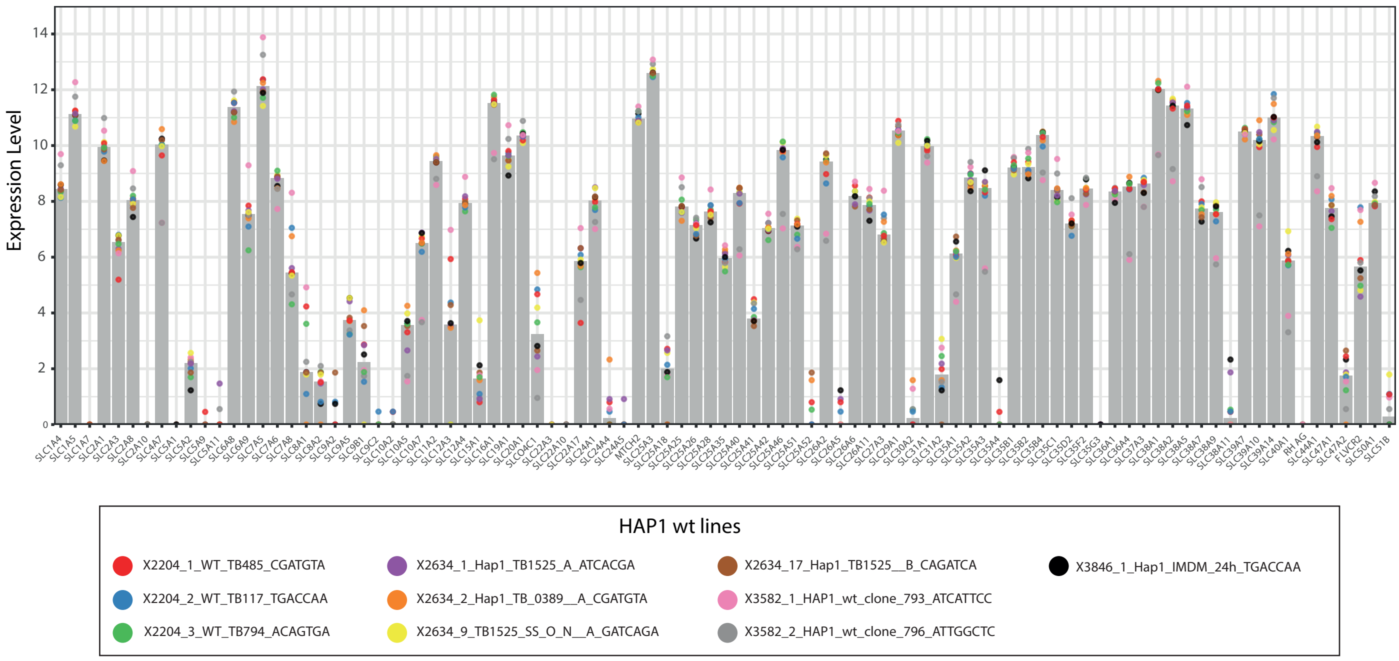


d

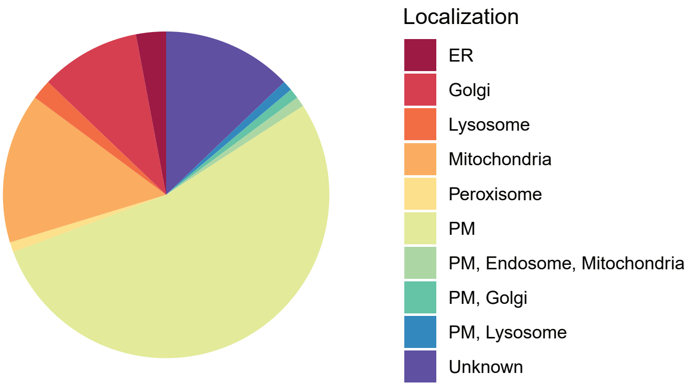

### Supplementary Figure 3

a

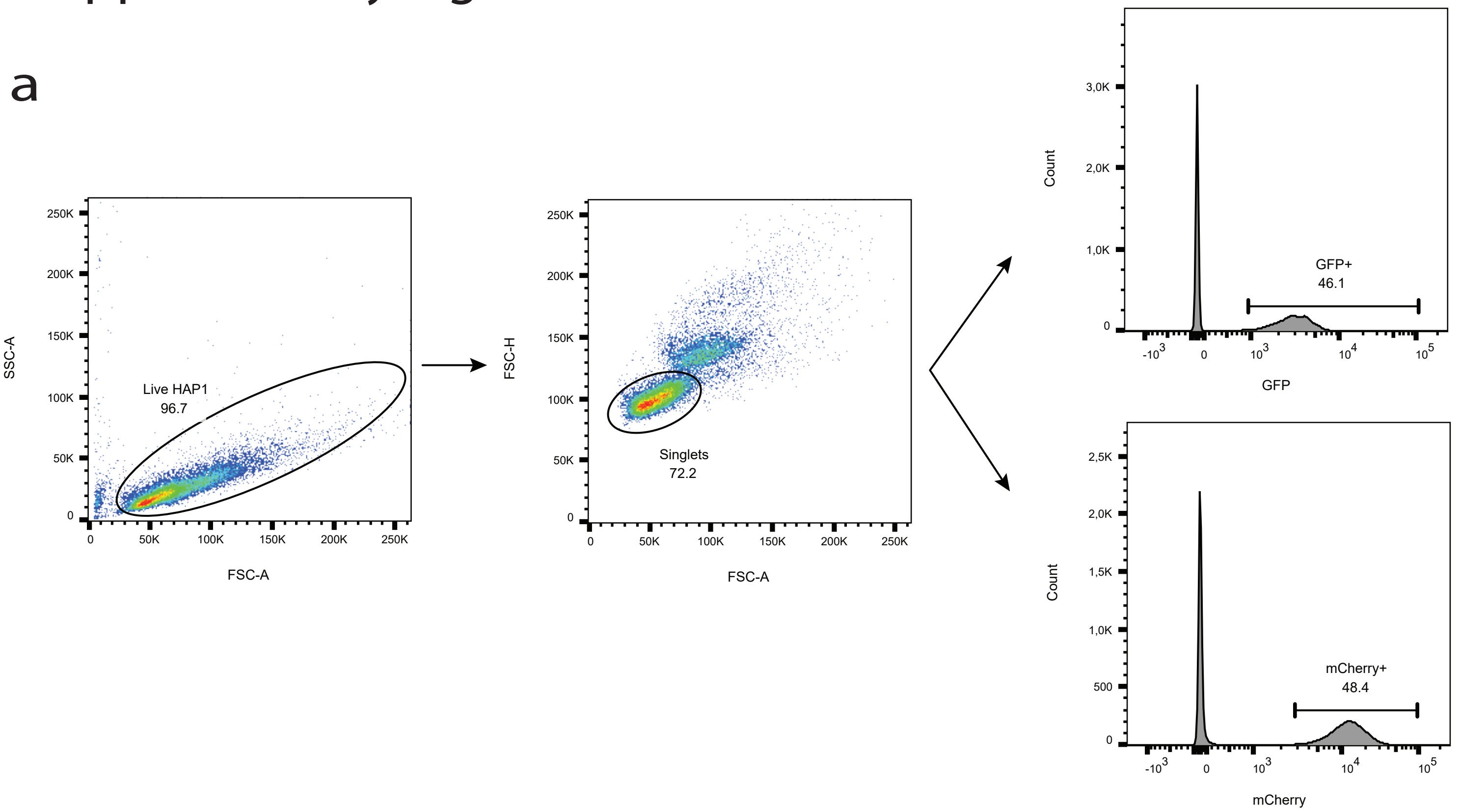

### Supplementary Figure 4

a

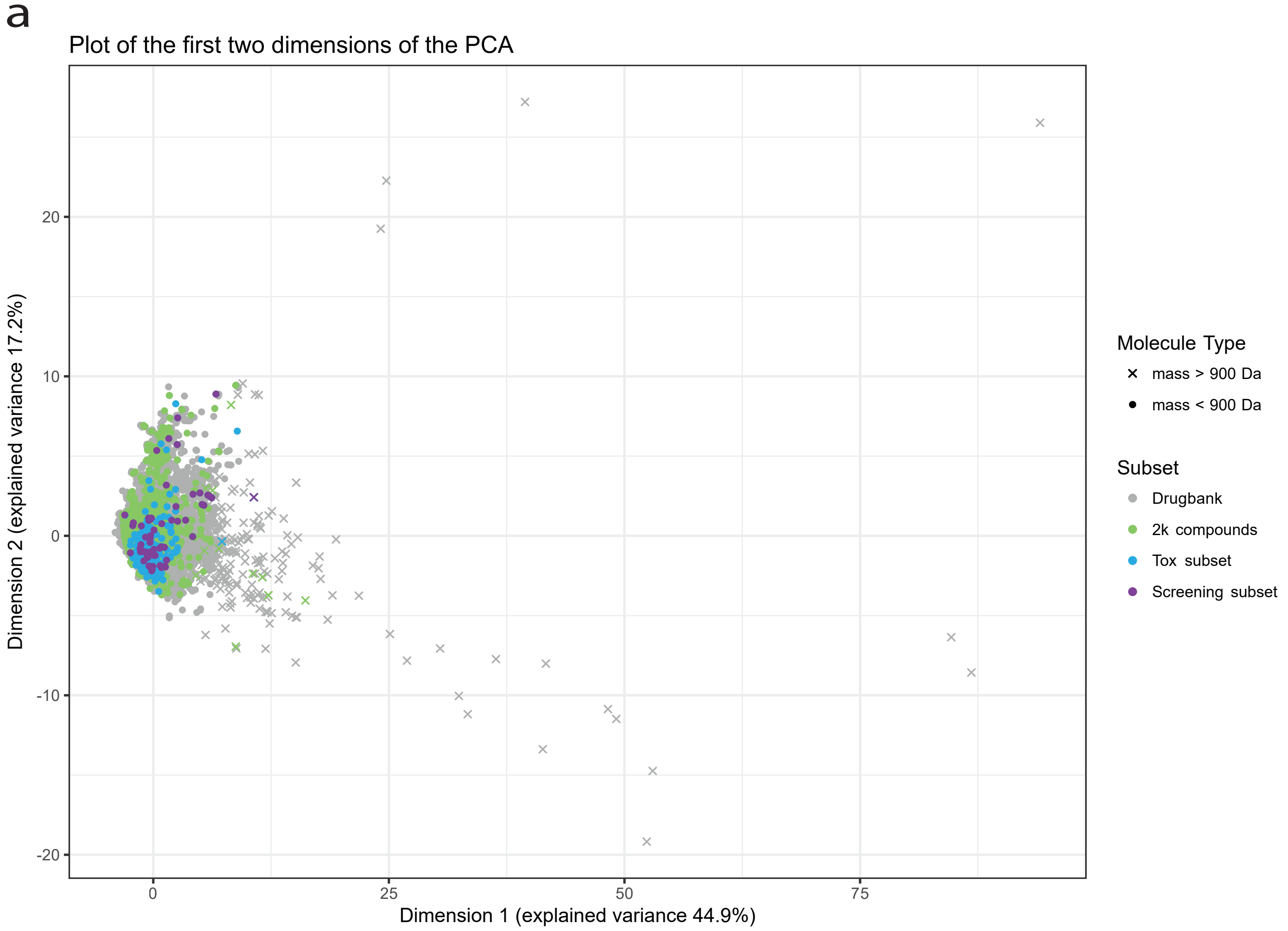


b

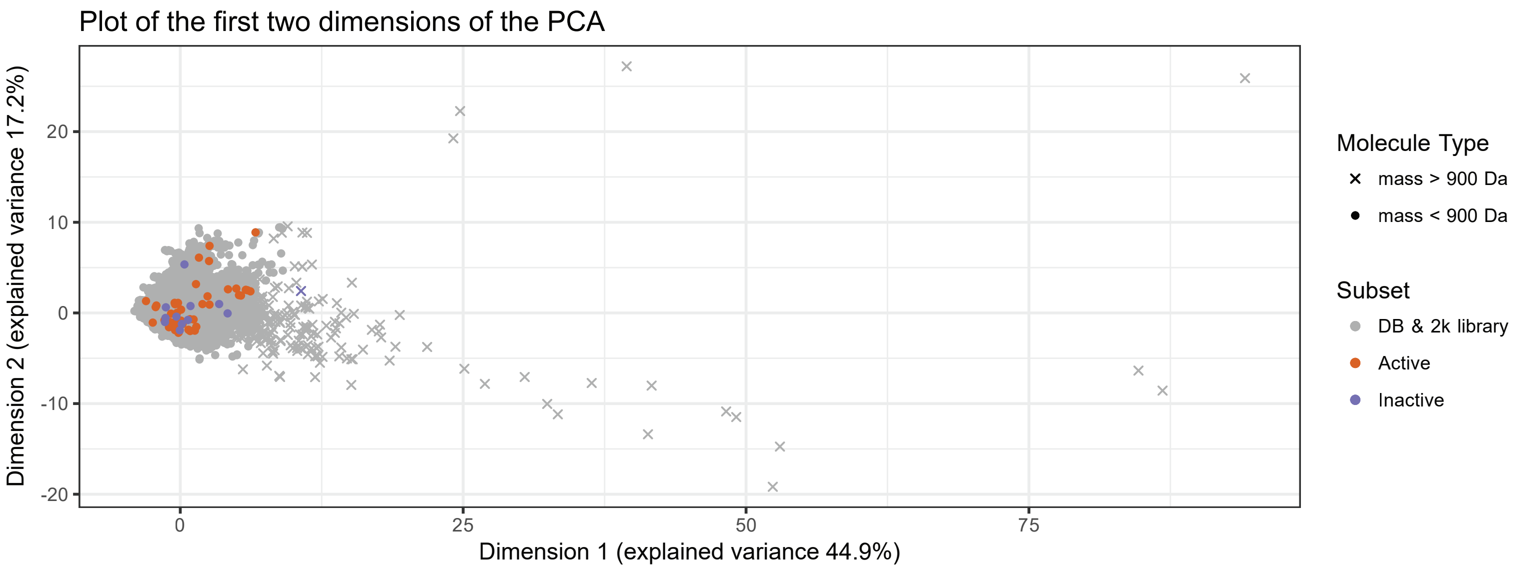


c

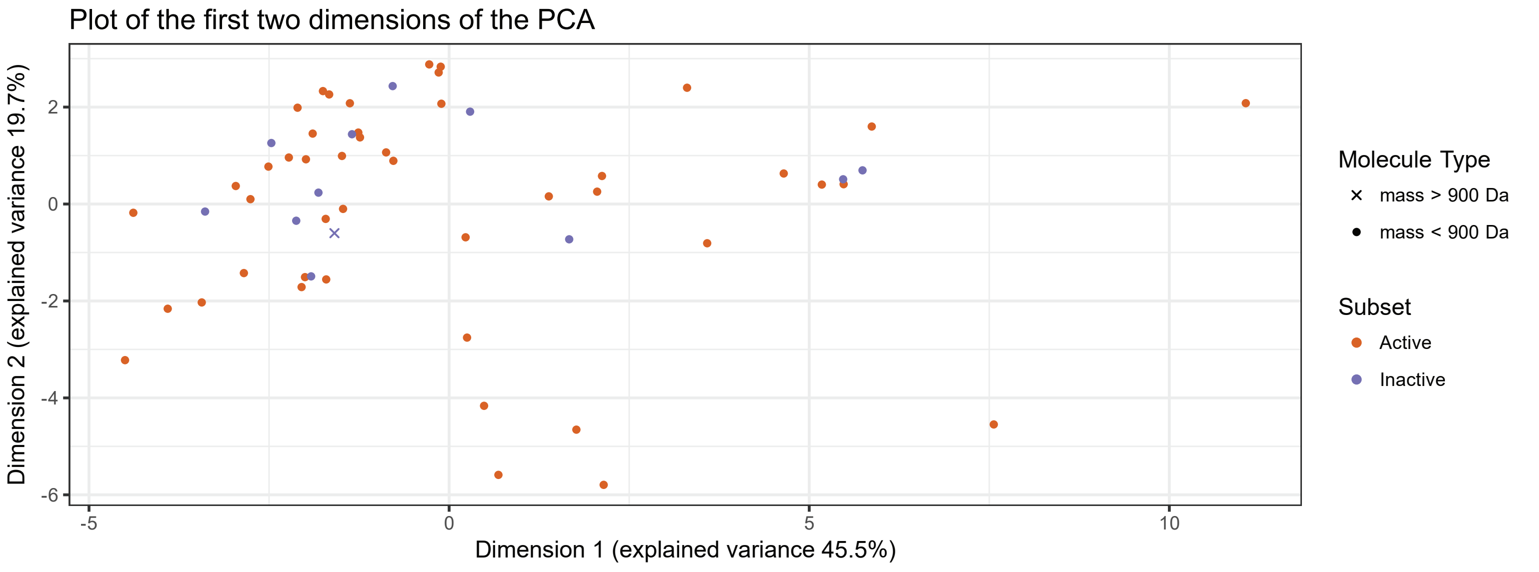

### Supplementary Figure 5

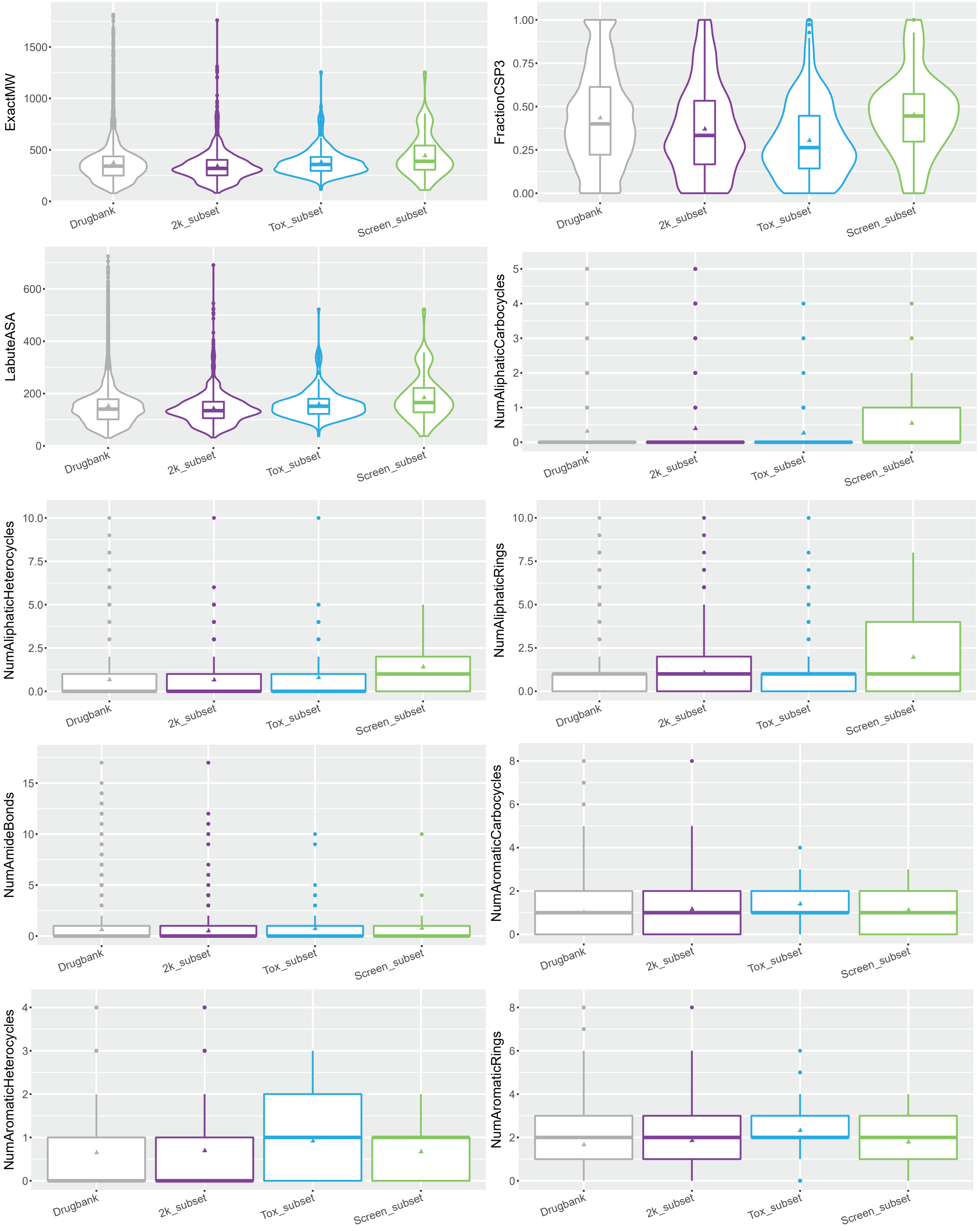

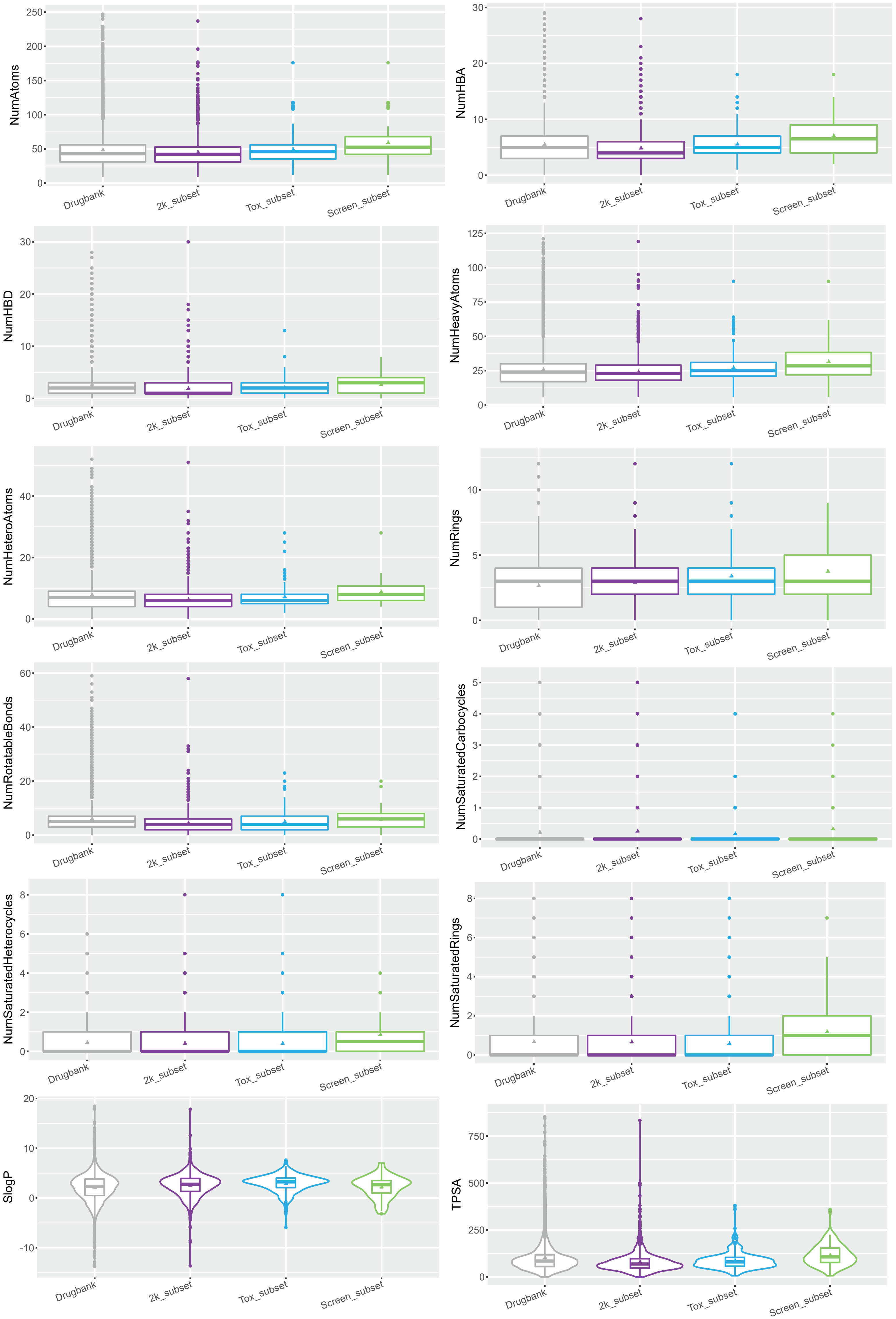

NumAtoms
0
50
100
150
200
250
NumHBA
0
10
20
30
NumHBD
0
10
20
30
NumHeavyAtoms
0
25
50
75
100
125
NumHeteroAtoms
0
20
40
NumRings
0
5
10
NumRotatableBonds
0
20
40
60
NumSaturatedCarbocycles
0
1
2
3
4
5
NumSaturatedHeterocycles
0
2
4
6
8
NumSaturatedRings
0
2
4
6
8
SlogP
-10
0
10
20
TPSA
0
250
500
750
Drugbank
2k_subset
Tox_subset
Screen_subset

**Supplemental Table 1**. Targeted SLCs and positive controls (~6 sgRNAs per gene)

| SLC genes and pseudogenes | | | | | | | | | Positive controls |
|---|---|---|---|---|---|---|---|---|---|
| SLC1A1 | SLC6A1 | *SLC9A7P1* | SLC16A10 | SLC22A12 | SLC25A26 | SLC27A6 | SLC35F2 | SLC41A1 | CEP85 |
| SLC1A2 | SLC6A2 | SLC9A8 | SLC16A11 | SLC22A13 | SLC25A27 | SLC28A1 | SLC35F3 | SLC41A2 | CTCF |
| SLC1A3 | SLC6A3 | SLC9A9 | SLC16A12 | SLC22A14 | SLC25A28 | SLC28A2 | SLC35F4 | SLC41A3 | CTNNBL1 |
| SLC1A4 | SLC6A4 | SLC9B1 | SLC16A13 | SLC22A15 | SLC25A29 | SLC28A3 | SLC35F5 | RHAG | DYRK1A |
| SLC1A5 | SLC6A5 | SLC9B2 | SLC16A14 | SLC22A16 | SLC25A30 | SLC29A1 | SLC35F6 | RHBG | KANSL1 |
| SLC1A6 | SLC6A6 | SLC9C1 | SLC17A1 | SLC22A17 | SLC25A31 | SLC29A2 | SLC35G1 | RHCG | MED26 |
| SLC1A7 | SLC6A7 | SLC9C2 | SLC17A2 | SLC22A18 | SLC25A32 | SLC29A3 | SLC35G2 | SLC43A1 | NBAS |
| SLC2A1 | SLC6A8 | SLC10A1 | SLC17A3 | SLC22A20 | SLC25A33 | SLC29A4 | SLC35G3 | SLC43A2 | NFYC |
| SLC2A2 | SLC6A9 | SLC10A2 | SLC17A4 | SLC22A23 | SLC25A34 | SLC30A1 | SLC35G4 | SLC43A3 | NIPBL |
| SLC2A3 | SLC6A11 | SLC10A3 | SLC17A5 | SLC22A24 | SLC25A35 | SLC30A2 | SLC35G5 | SLC44A1 | NUP153 |
| *SLC2A3P1* | SLC6A12 | SLC10A4 | SLC17A6 | SLC22A25 | SLC25A36 | SLC30A3 | SLC35G6 | SLC44A2 | PAFAH1B1 |
| SLC2A4 | SLC6A13 | SLC10A5 | SLC17A7 | SLC22A31 | SLC25A37 | SLC30A4 | SLC36A1 | SLC44A3 | RCC1 |
| SLC2A5 | SLC6A14 | SLC10A6 | SLC17A8 | SLC23A1 | SLC25A38 | SLC30A5 | SLC36A2 | SLC44A4 | RPL13A |
| SLC2A6 | SLC6A15 | SLC10A7 | SLC17A9 | SLC23A2 | SLC25A39 | SLC30A6 | SLC36A3 | SLC44A5 | RPTOR |
| SLC2A7 | SLC6A16 | SLC11A1 | SLC18A1 | SLC23A3 | SLC25A40 | SLC30A7 | SLC36A4 | SLC45A1 | SAE1 |
| SLC2A8 | SLC6A17 | SLC11A2 | SLC18A2 | SLC24A1 | SLC25A41 | SLC30A8 | SLC37A1 | SLC45A2 | SRBD1 |
| SLC2A9 | SLC6A18 | SLC12A1 | SLC18A3 | SLC24A2 | SLC25A42 | SLC30A9 | SLC37A2 | SLC45A3 | U2AF2 |
| SLC2A10 | SLC6A19 | SLC12A2 | SLC18B1 | SLC24A3 | SLC25A43 | SLC30A10 | SLC37A3 | SLC45A4 | URI1 |
| SLC2A11 | SLC6A20 | SLC12A3 | SLC19A1 | SLC24A4 | SLC25A44 | SLC31A1 | SLC37A4 | SLC46A1 | VMP1 |
| SLC2A12 | SLC7A1 | SLC12A4 | SLC19A2 | SLC24A5 | SLC25A45 | SLC31A2 | SLC38A1 | SLC46A2 | VPS13D |
| SLC2A13 | SLC7A2 | SLC12A5 | SLC19A3 | MTCH1 | SLC25A46 | SLC32A1 | SLC38A2 | SLC46A3 | |
| SLC2A14 | SLC7A3 | SLC12A6 | SLC20A1 | MTCH2 | SLC25A47 | SLC33A1 | SLC38A3 | SLC47A1 | |
| SLC3A1 | SLC7A4 | SLC12A7 | SLC20A2 | SLC25A1 | SLC25A48 | SLC34A1 | SLC38A4 | SLC47A2 | |
| SLC3A2 | SLC7A5 | SLC12A8 | SLCO1A2 | *SLC25A1P5* | SLC25A51 | SLC34A2 | SLC38A5 | SLC48A1 | |
| SLC4A2 | *SLC7A5P1* | SLC12A9 | SLCO1B1 | SLC25A2 | SLC25A52 | SLC34A3 | SLC38A6 | DIRC2 | |
| SLC4A3 | *SLC7A5P2* | SLC13A1 | SLCO1B3 | SLC25A3 | SLC25A53 | SLC35A1 | SLC38A7 | FLVCR1 | |
| SLC4A4 | SLC7A6 | SLC13A2 | SLCO1C1 | SLC25A4 | UCP1 | SLC35A2 | SLC38A8 | FLVCR2 | |
| SLC4A5 | SLC7A7 | SLC13A3 | SLCO2A1 | SLC25A5 | UCP2 | SLC35A3 | SLC38A9 | MFSD7 | |
| SLC4A7 | SLC7A8 | SLC13A4 | SLCO2B1 | SLC25A6 | UCP3 | SLC35A4 | SLC38A10 | SLC50A1 | |
| SLC4A8 | SLC7A9 | SLC13A5 | SLCO3A1 | SLC25A10 | SLC26A1 | SLC35A5 | SLC38A11 | SLC51A | |
| SLC4A9 | SLC7A10 | SLC14A1 | SLCO4A1 | SLC25A11 | SLC26A2 | SLC35B1 | SLC39A1 | SLC51B | |
| SLC4A10 | SLC7A11 | SLC14A2 | SLCO4C1 | SLC25A12 | SLC26A3 | SLC35B2 | SLC39A2 | SLC52A1 | |
| SLC4A11 | SLC7A13 | SLC15A1 | SLCO5A1 | SLC25A13 | SLC26A4 | SLC35B3 | SLC39A3 | SLC52A2 | |
| SLC5A1 | SLC7A14 | SLC15A2 | SLCO6A1 | SLC25A14 | SLC26A5 | SLC35B4 | SLC39A4 | SLC52A3 | |
| SLC5A2 | SLC8A1 | SLC15A3 | SLC22A1 | SLC25A15 | SLC26A6 | SLC35C1 | SLC39A5 | | |
| SLC5A3 | SLC8A2 | SLC15A4 | SLC22A2 | SLC25A16 | SLC26A7 | SLC35C2 | SLC39A6 | | |
| SLC5A4 | SLC8A3 | SLC16A1 | SLC22A3 | SLC25A17 | SLC26A8 | SLC35D1 | SLC39A7 | | |
| SLC5A5 | SLC8B1 | SLC16A2 | SLC22A4 | SLC25A18 | SLC26A9 | SLC35D2 | SLC39A8 | | |
| SLC5A6 | SLC9A1 | SLC16A3 | SLC22A5 | SLC25A19 | SLC26A10 | SLC35D3 | SLC39A9 | | |
| SLC5A7 | SLC9A2 | SLC16A4 | SLC22A6 | SLC25A20 | SLC26A11 | SLC35E1 | SLC39A10 | | |
| SLC5A8 | SLC9A3 | SLC16A5 | SLC22A7 | SLC25A21 | SLC27A1 | *SLC35E1P1* | SLC39A11 | | |
| SLC5A9 | SLC9A4 | SLC16A6 | SLC22A8 | SLC25A22 | SLC27A2 | SLC35E2B | SLC39A12 | | |
| SLC5A10 | SLC9A5 | SLC16A7 | SLC22A9 | SLC25A23 | SLC27A3 | SLC35E3 | SLC39A13 | | |
| SLC5A11 | SLC9A6 | SLC16A8 | SLC22A10 | SLC25A24 | SLC27A4 | SLC35E4 | SLC39A14 | | |
| SLC5A12 | SLC9A7 | SLC16A9 | SLC22A11 | SLC25A25 | SLC27A5 | SLC35F1 | SLC40A1 | | |

**Supplemental Table 2**. Screened compounds.

| Class | Subclass | Name | Status* | IC50 in HAP1 cells (µM) |
|---|---|---|---|---|
| antineoplastic | purine analogs | 6-mercaptopurine | A | 3,176 |
| | nucleoside analogs | 5-azacitidine | A | 11,07 |
| | | clofarabine | A | 0,1129 |
| | | cytarabine | A | 0,3032 |
| | | decitabine | A | 0,7 |
| | | gemcitabine | A | 0,0075 |
| | antifolates | methotrexate | A | 0,1 |
| | | pralatrexate | A | 8,3 |
| | | raltitrexed | A | 0,03902 |
| | HDAC inhibitors | belinostat | A | 0,3215 |
| | | chidamide (tucidinostat) | I | 3,39 |
| | | entinostat | I | 1,95 |
| | | panobinostat | A | 0,01022 |
| | | pracinostat | I | 0,3094 |
| | | resminostat | I | 2,478 |
| | | romidepsin | A | 10,5 |
| | | vorinostat | A | 2,894 |
| | microtubule inhibitors (destabilizing) | vinblastine | A | 9,8 |
| | | vincristine | A | 28 |
| | | vindesine | A | 0,00543 |
| | | vinorelbine | A | 0,1012 |
| | microtubule inhibitors (stabilizing) | docetaxel | A | 0,02 |
| | | paclitaxel | A | 6,3 |
| | proteasome inhibitors | bortezomib | A | 0,00649 |
| | | carfilzomib | A | 3,6 |
| | RTK inhibitors | crizotinib | A | 3,6 |
| | | ponatinib | A | 0,46 |
| | | sunitinib | A | 2,891 |
| | topoisomerase I inhibitors | topotecan | A | 0,006 |
| | | irinotecan | A | 0,7631 |

*based on DrugBank: A (approved), A, W (approved, withdrawn), I (investigational), E (experimental)

**Supplemental Table 2 (cont)**. Screened compounds.

| Class | Subclass | Name | Status* | IC50 in HAP1 cells (μM) |
|---|---|---|---|---|
| antineoplastic | topoisomerase II inhibitors | doxorubicin | A | 0,007 |
| | | epirubicin | A | 0,017 |
| | | etoposide | A | 0,238 |
| | | idarubicin | A | 0.022 |
| | | mitoxantrone | A | 0,007 |
| | protein translation inhibitors | homoharringtonine (omacetaxine mepesuccinate) | A | 0,025 |
| | transcription inhibitors | dactinomycin | A | 0,039 |
| | alkylating | cisplatin | A | 1 |
| | | methyl methanesulfonate | A | 24 |
| | | mitomycin C | A | 15 |
| | | temozolomide | A | 5 |
| | other | triptolide | I | 0,005 |
| antiparasitic | antimalarial | artesunate | A | 1,786 |
| | | dihydroartemisinin (artenimol) | I | 6,3 |
| | | mefloquine | A | 7,914 |
| | antihelmintic | albendazole | A | 0,452 |
| | antiprotozoal | pentamidine | A | 6,206 |
| antiarrhythmic | type III: K-channel blocker | amiodarone | A | ~10 |
| | | dronedarone | A | 6,3 |
| | type V | digitoxin | A | 0,001 |
| antihypertensive | Ca-channel blocker | nisoldipine | A | 9,8 |
| anti-inflammatory | NSAID | oxyphenbutazone | A, W | ~10 |
| immunosuppressant | | mycophenolic acid | A | 0,934 |
| hypolipidemic | HMG-CoA reductase inhibitor | cerivastatin | A, W | 0,15 |
| antipasmodic | | chlorzoxazone | A | ~10 |
| | | metaxalone | A | ~10 |
| prokinetic | serotonin agonist | tegaserod | A, W | 5,856 |
| mineralocorticoid | | desoxycorticosterone pivalate | E | ~10 |
| uricosuric | | sulfinpyrazone | A | 10 |
| alcohol deterrent | | disulfiram | A | 28 |

*based on DrugBank: A (approved), A, W (approved, withdrawn), I (investigational), E (experimental)

**Supplemental Table 3**. Descriptors used in the chemical space analysis

| Descriptor name | Description |
|---|---|
| SlogP | Smarts LogP, Octanol Water Partition Coefficient |
| LabuteASA | Labute's Approximate Surface Area, approximated surface area of a molecule (J. Mol. Graph. Mod. 18, 464-77 (2000)) |
| TPSA | Total Polar surface area |
| ExactMW | Molecular weight |
| NumRotatableBonds | Number of rotatable bonds |
| NumHBD | Number of hydrogen bond donors |
| NumHBA | Number of hydrogen bond acceptors |
| NumAmideBonds | Number of amide bonds |
| NumHeteroAtoms | Number of hetero atoms |
| NumHeavyAtoms | Number of heavy atoms |
| NumAtoms | Number of atoms |
| NumRings | Number of rings |
| NumAromaticRings | Number of aromatic rings |
| NumSaturatedRings | Number of saturated rings |
| NumAliphaticRings | Number of aliphatic rings |
| NumAromaticHeterocycles | Number of aromatic heterocycles |
| NumSaturatedHeterocycles | Number of saturated heterocycles |
| NumAliphaticHeterocycles | Number of aliphatic heterocycles |
| NumAromaticCarbocycles | Number of aromatic carbocycles |
| NumSaturatedCarbocycles | Number of saturated carbocycles |
| NumAliphaticCarbocycles | Number of aliphatic carbocycles |
| FractionCSP3 | Fraction of sp3 hybridized Carbons |

**Supplemental Table 4**. Selected SLC-drug interactions for validation

| *Gene* | *Drug* | *Concentration (IC50)* |
|---|---|---|
| **SLC1A5** | Mitoxantrone | 3 |
| | Vinorelbine | 3 |
| | Homoharringtonine | 1 |
| | Panobinostat | 3 |
| | Entinostat | 3 |
| **SLC11A2** | Artesunate | 3 |
| | Dihydroarthemisinin | 3 |
| **SLC16A1** | Artesunate | 3 |
| | Dihydroarthemisinin | 3 |
| | Nisoldipine | 3 |
| **SLC19A1** | Pralatraxate | 3 |
| | Raltitrexed | 3 |
| | Pentamidine | 3 |
| | Methotrexate | 3 |
| **SLC20A1** | Pentamidine | 3 |
| | Methotrexate | 3 |
| **SLC25A3** | Decitabine | 3 |
| | Cytarabine | 1 |
| **MTCH2** | Decitabine | 3 |
| | Cytarabine | 3 |
| | Nisoldipine | 3 |
| | Belinostat | 3 |
| **SLC29A1** | Gemcitabine | 1 |
| | Topotecan | 1 |
| | Decitabine | 3 |
| | Cytarabine | 3 |
| | 5-azacytidine | 3 |
| **SLC35A1** | Sulfinpyrazone | 3 |
| | Digitoxin | 1 |
| **SLC35A2** | Cisplatin | 1 |
| | 5-azacytidine | 1 |
| **SLC38A5** | Cisplatin | 1 |
| **SLC47A1** | Mitoxantrone | 3 |
| **SLC47A2** | Mitoxantrone | 3 |

**Supplemental Table 5**. sgRNAs used to generate MCA cell lines and corresponding editing efficiencies as assessed by Tide-seq.

| Gene | KO | %editing | sgRNA |
|---|---|---|---|
| SLC1A5 | 1 | 83,8 | GCCGCTGATGATGAAGTGCG |
| | 2 | 67,8 | CAGCGCCACACCAAAGACGA |
| SLC11A2 | 1 | 39,1 | ATCAGCCACAGGATGACTCG |
| | 2 | 52,1 | ATGAGAACGCCACCCCACAG |
| SLC16A1 | 1 | 54 | ACAGACGTATAGTTGCTGTA |
| | 2 | 42,8 | TATCCATGACACTTCGCTGG |
| SLC19A1 | 1 | 40,3 | GGCCCGACAAGAACTTCACG |
| | 2 | 42,8 | CGACTACCTGCGCTACACGC |
| SLC20A1 | 1 | 51,1 | TTGGCACGGAATGAATCCAG |
| | 2 | 42,4 | CAGGCCGGAATCCTTATGCA |
| SLC25A3 | 1 | NA | TTCAACAGTACGTTCAAAGC |
| | 2 | 57 | TCTGATCTCACCTTCCACGG |
| MTCH2 | 1 | 35,5 | ACATTGCCAGTATCGATGGG |
| | 2 | 10,7 | AGCACTTTCACGTACATGAG |
| SLC29A1 | 1 | 49,2 | GCTCAAGCTTGAAGGACCCG |
| | 2 | 40,7 | GCTCAAGCTTGAAGGACCCG |
| SLC35A1 | 1 | 32,6 | TGAACAGCATACACTAACGA |
| | 2 | 19,2 | ACACGGAATCTTCAACTGGT |
| SLC35A2 | 1 | 29,6 | TAGAGATGGCAACATACTGG |
| | 2 | 13,5 | CTACGCCCGCACGTTGCCAG |
| SLC38A5 | 1 | 44,5 | CTATGCCATGGCCCACACGG |
| | 2 | 32,4 | TATCGGCACC TTCCTGTACA |
| SLC47A1 | 1 | 17,7 | AGCCAGAACCTGAAGCACGT |
| | 2 | 42,9 | GCAACTCCAGTTACGATCTG |
| SLC47A2 | 1 | 36,5 | GGCATCGGTGACCCTCGCGG |
| | 2 | 33,4 | GCTGGCATCG GTGACCCTCG |
